# Migrating mesoderm cells self-organize into a dynamic meshwork structure during chick gastrulation

**DOI:** 10.1101/2022.09.08.507227

**Authors:** Yukiko Nakaya, Mitsusuke Tarama, Sohei Tasaki, Ayako Isomura, Tatsuo Shibata

**Author notes:** **Corresponding author**, Tatsuo Shibata.

## Abstract

Migration of cell populations is a fundamental process in morphogenesis and disease. The mechanisms of collective cell migration of epithelial cell populations have been well studied. It remains unclear, however, how the highly motile mesenchymal cells, which migrate extensively throughout the embryo, are connected with each other and coordinated as a collective. During gastrulation in chick embryos, the mesoderm cells, that are formed by an epithelial-to-mesenchymal transition (EMT), migrate in the 3D space between ectoderm and endoderm of the embryo. Using live imaging and quantitative analysis, such as topological data analysis (TDA), we found that the mesoderm cells undergo a novel form of collective migration, in which they form a meshwork structure while moving away from the primitive streak. This meshwork is supported by N-cadherin-mediated cell-cell adhesion, which undergoes rapid reorganization. Overexpressing a mutant form of N-cadherin decreases the speed of tissue progression and the directionality of the collective cell movement, whereas the speed of individual cells remains unchanged. To investigate how this meshwork arises and how it contributes to the cell movement, we utilized an agent-based theoretical model, showing that cell elongation, cell-cell adhesion, and cell density are the key parameters for the meshwork formation. These data provide novel insights into how a supracellular structure of migrating mesenchymal cells forms and how it facilitates efficient migration during early mesoderm formation.

## Introduction

Collective behaviors of migrating cells are fundamental in processes of morphogenesis of tissues and organ, wound healing, and tumor metastasis. Such collective cell migration is typically found in epithelial tissues, in which the motile ability can be acquired by bringing multiple cells together into one group via intercellular adhesion (1–5). On the other hand, mesenchymal cells do not exhibit stable intercellular adhesion with surrounding cells, and they can migrate as individual cells. However, even for these mesenchymal cells, collective cell migration that exploits transient cell-cell adhesion is required for the morphogenesis in living organisms (1, 6, 7). Neural crest (NC) migration is one of the most studied model systems for mesenchymal collective migration, in which cells are gathered into characteristic chains or streams within a physically restricted environment (8). In *Xenopus* embryos, cranial NC streams emerge from the interaction with neighboring tissue placode, where transient cell-cell interactions called contact inhibition of locomotion confer the supracellular polarity to determine the orientation of movement as a group (9, 10). Whereas the mechanism of this streaming migration is relatively well investigated (11), it is still largely unknown how mesenchymal cells that are not tightly confined, such as those in mesoderm, move toward their destination and how their transient cell-cell adhesions contribute to it.

Mesoderm is a germ layer consisting of mesenchymal cells, and it forms during gastrulation. In the case of chick embryos, as the primitive streak is being formed, cells in the superficial layer (epiblast) move toward the primitive streak. Most of the mesoderm cells are formed by the convergence of these epithelial-shaped epiblast cells to the primitive streak, which subsequently undergo the epithelial-to-mesenchymal transition (EMT). These mesoderm cells ingress adopting an irregular mesenchymal morphology and acquire high motility (12–14). Then, the mesoderm cells move away from the primitive streak at various anterior-posterior positions, in the three-dimensional (3D) space between the epiblast and endoderm (Figure 1A) (movie: https://www.sdbcore.org/object?ObjectID=358) (12–18). Previous reports suggested that the mesoderm cells migrate at high density and their long-range migration pathway was controlled by a balance between chemo-repulsion mediated by FGF8 secreted from the primitive streak and chemo-attraction mediated by FGF4 secreted from the head-process and notochord (3, 19). It is not clear, however, whether they are essentially solitary cells following the same cues while occasionally contacting each other or whether collective effects are essential for the mesoderm migration. If the latter is the case, questions arise as to what kind of cell-cell interactions give rise to the collective property, and what spatial structure emerges beyond the single cellular scale when cells migrate collectively.

**Figure1.**
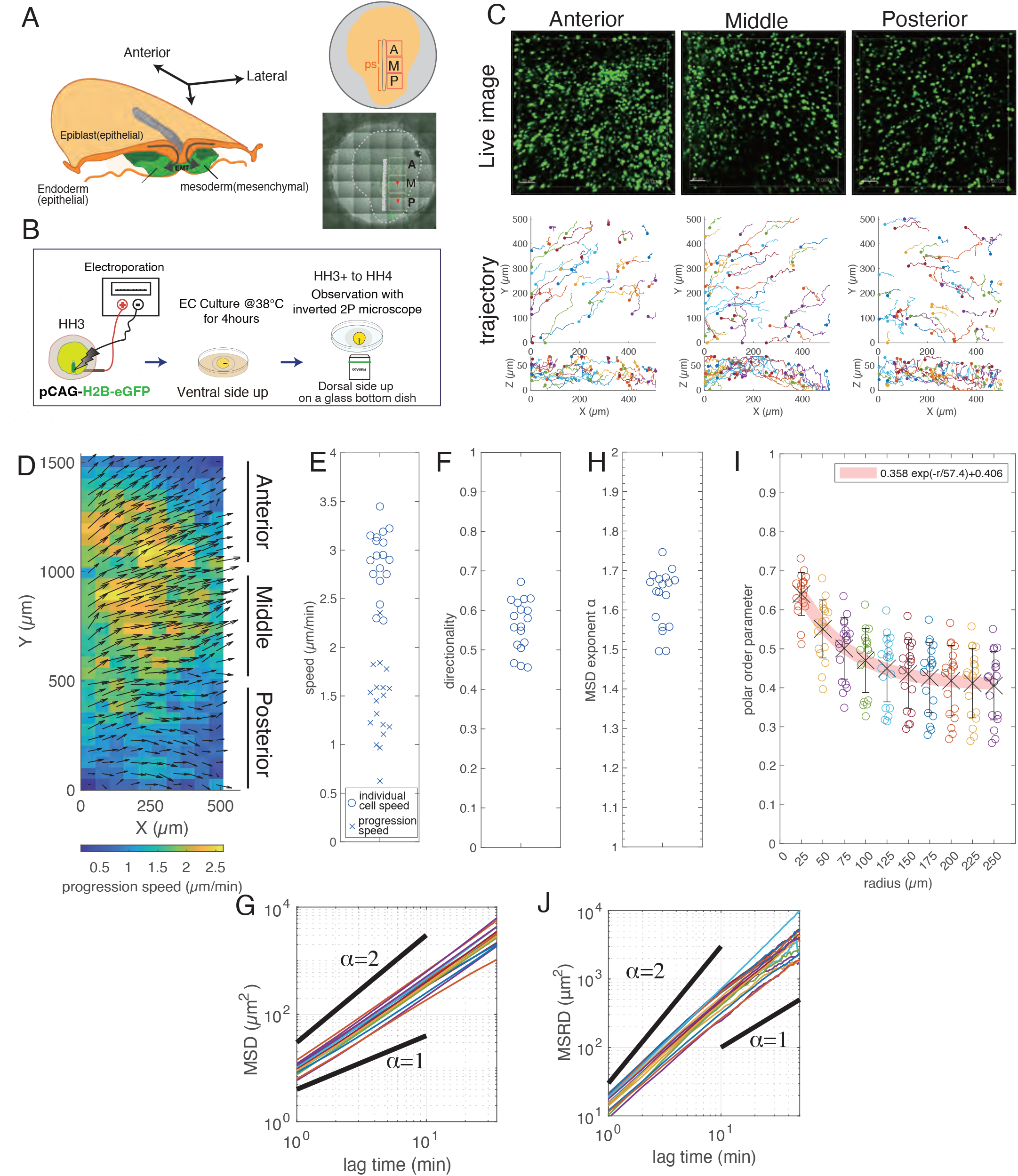
Mesoderm cells move collectively during gastrulation. (A) Schematic diagram of the chicken embryo at stage HH3. The observation regions are marked by the square boxes in the right panels (A: Anterior, M: Middle, P: Posterior). (B) Experimental procedure. DNA encoding H2B-eGFP was introduced into the cells in the primitive streak at stage HH3 by electroporation. After several hours of incubation, the position of the labeled nuclei was recorded using a multi-photon microscope. (C) Examples of the obtained images of the mesoderm cells expressing H2B-eGFP (upper panels) and reconstructed 3D trajectories (bottom panels). The x, y, and z axes correspond to the mediolateral, anterior-posterior, and dorsoventral axes, respectively. Scale bar: 50μm. (D) Spatial distribution of progression velocity (arrows) and progression speed (color). (E) Individual cell speed (o) and progression speed (x), and (F) directionality. Each data point of the individual cell speed and the directionality represents the average over the cells and that of the progression speed is the average over the subareas in each region of the 6 embryos. (G) Mean squared displacement (MSD). Each line corresponds to the MSD in each region of the 6 embryos. (H) Exponent of the MSD plotted in (G). (I) Polar order parameter *φ* plotted against the radius of the measurement area. The polar order parameter calculated for the cells in the circular areas of a given radius at each time was averaged over the areas and time in each region of the 6 embryos (the crosses and error bars). (J) Mean squared relative distance (MSRD). Each line represents the MSRD in each region of the 6 embryos. On a time scale larger than about 10 min, the exponent of the MSRD becomes 1. The numbers of cells analyzed are N=1525 (A), 1112(M), 791(P) (embryo 1), 371(A), 416 (M), 235 (P) (embryo 2), 398 (A), 388 (M), 316 (P) (embryo 3), 230 (A), 296 (M), 175 (P) (embryo 4), 1386 (A), 1283 (M), 964 (P) (embryo 5), 1040 (A), 1102 (M), 496 (P) (embryo 6).

In this study, we investigated the cellular mechanisms underlying mesoderm cell migration in chicks. Our quantitative data analysis including large-scale cell tracking and topological data analysis (TDA) on high-resolution microscopy images showed that the mesoderm cells migrate collectively, and they are connected to each other via cadherin-mediated cell-cell adhesion and form a meshwork structure that exhibits a continual and rapid reorganization. This dynamic meshwork structure was reproduced by a theoretical model, which demonstrates that the morphology of the cells, the strength of cell-cell adhesion, and the cell density are key parameters for the meshwork formation.

## Results

### Mesoderm cells move collectively with frequent changes in their relative position

We first investigated how mesoderm cells migrate in the early gastrulating stage of chick embryos. To this end, we electroporated a plasmid encoding H2B-eGFP into the mesoderm precursor cells located at the primitive streak of stage HH3 embryos (20), and then the embryos were cultured *ex vivo* for several hours (21). To follow the behavior of electroporated cells, we obtained 3D images (scan area:500μm x 500μm, scan depth:50μm) of the mesoderm at three different streak levels along the primitive streak, at 1 min intervals, for 2 h using an inverted two-photon microscope with 25x WMP/0.25 NA objective (Figures 1A and 1B, and Video S1). We note that at this developmental stage (HH3) the entire length of the primitive streak is less than 2mm. We also note that although the tissue scale extension along the anterior-posterior axis becomes more prominent after HH4+, the cell trajectories at this stage (HH3) reflect the cell migration in the environment of mesoderm region without the tissue-scale deformation.

From the position of the nuclei, we reconstructed the 3D trajectories of mesoderm cells (Figure 1C and Video S1), where the x and y axes correspond to the mediolateral and anterior-posterior (AP) axes, respectively. Most of these 3D trajectories were of mesoderm cells, but there might have been a few endodermal and epiblast cells. In all three regions, the mesoderm cells exhibited a spreading behavior with maximal cell displacement in particular directions (Figures 1C, Anterior, Middle, and 1D). In the anterior and middle regions, the mesoderm cells tended to move in the anterior-lateral direction (Figures 1C Anterior, Middle and 1D). In contrast, the mesoderm cells in the posterior region tended to move toward the lateral direction (Figures 1C, Posterior and 1D). These migration directions were consistent with previous reports (17, 19). Now we quantitatively investigate the migration of individual mesoderm cells and how their migration are coordinated among neighboring cells.

The trajectory of the individual cells showed that the mesoderm cells frequently changed their direction of motion, and some cells move even toward the primitive streak transiently (Figure 1C). The z-position of the mesoderm cells also changed frequently (Figure 1C). In fact, the distribution of the difference between the maximum and minimum z-positions of individual trajectories in 60 min showed a peak around 20 μm with a tail that extended more than 40 μm (Figure S1A), indicating that most cells change their z-position by more than one to two cell lengths, given that the typical cell length is 10 to 20 μm. Thus, the mesoderm cells frequently change their direction of motion in 3D.

We first quantitatively characterize this migration behavior of individual cells for trajectories obtained from six embryos (in total, 6 × 3 = 18 samples) (see SI Methods). The mean values of individual cell speed for each sample were distributed from about 2.3 to 3.5 *μm/min* (Figure 1E) (for the speed of individual cell, see Figure S1B). The directionality defined by the ratio of the start-to-end distance to the total path length measures how persistent the trajectories are (SI Methods). The directionality for the trajectories of 20 min length in 18 samples was smaller than unity and was distributed from about 0.45 to 0.7 (Figure 1F). We then performed a mean squared displacement (MSD) analysis to evaluate the randomness of the individual mesoderm migration (SI Methods). The MSD were proportional to *t^α^*, where *α* was distributed from about 1.5 to 1.75 (Figure. 1G and H). This result means that the mesoderm cell motion in all regions is in the regime between random walk (*α* = 1) and ballistic motion (*α* = 2) (22). We also confirmed that the MSD exponent and the directionality showed a good positive correlation (Figure S1C). To further verify the persistence of the migration direction, we calculated the autocorrelation function (ACF) of the velocity of individual cells (Figure S1D). The ACF indicated that the correlation decreased below 0.5 in 2 min for all 18 samples, indicating that the migration direction shows random fluctuations. For longer time scale, the ACF gradually decreased, but did not converge to zero, indicating that the motion was biased slightly to a particular direction in each region. Thus, the motion of mesoderm cell is considered as a biased random walk. We note that these migration characteristics were not significantly different between the three regions, anterior, middle, and posterior.

We next elucidate the coordination of cell migrations among neighboring cells by analyzing the collective order of the mesoderm cell motions. To this end, we measured the polar order parameter *φ* of the direction of cell motion that quantifies the instantaneous directional alignment among cells (SI Methods) (23). When all cells move in the same direction, *φ* is unity, whereas *φ* is zero when the direction of the cell migration is completely random. First, we calculated *φ* for cells within a circular region of radius 25 *μm* in the xy-coordinate. The value of *φ* was distributed from 0.5 to 0.75 with a mean value of 0.64, indicating that the migration direction is well aligned among the neighboring cells (Figure 1I), despite of the randomness in the individual cell motion. To investigate how far the cell migration direction is aligned, we measured the polar order parameters for different radius of the measurement circle. As the radius of the measurement circle increases, the polar order parameter decreases exponentially with a characteristic decay length of 57 *μm*, beyond which the average of *φ* is less than 0.5. This result indicates that the mesoderm cells within a length scale of about 60 *μm* migrate collectively. In the longer length scale, the migration of the mesoderm cells becomes less coordinated due to the fluctuation in the migration direction as the ACF of the velocity indicates.

To study whether the mesoderm cells within this characteristic length move together, we measured the mean squared relative distance (MSRD) (SI Methods), which quantifies the temporal change of the relative distance between two cells. The MSRD curve increases with time, but there is a threshold time about 10 min (Figure 1J) beyond which the increase of the MSRD slows down. In this time scale, the two cells initially in contact move away from each other to about 25 μm. Beyond the threshold time, the MSRD increases linearly in time. This indicates that the migrating mesoderm cells do not form a tight cluster, changing their relative positions frequently. It also suggests that there is sufficient extracellular space where the cells can easily change their relative positions in the mesoderm tissue.

We finally measured the progression speed of mesoderm for 18 samples, which was obtained as the time average of the average velocity of cells in small regions (50 μm x 50 μm) (Figure 1D color code). The mean of the progression speed was distributed from about 0.5 to 2 μm/min (Figure 1E), which was almost the half of the individual cells speed (Figures 1E and S1B). This difference can be explained by considering the randomness of cell migrations and the value of the polar order parameter of this length scale.

### Mesoderm cells form meshwork structure

If there is sufficient extracellular space for cells to change their relative position frequently, how do they distribute in 3D space between the epiblast and the endoderm? To explore this question, we prepared 3D reconstructed images of the mesoderm tissue at the mid-streak level in HH4 embryos (Figure 2A). From the projected image on the xy plane (Figure 2B, z-projection), we found no clear pattern in the distribution of cells in the primitive streak (dense region of nuclei stained with DAPI) and the mesoderm next to the streak, consistent with previous reports based on scanning electron microscopy (SEM) analysis (24, 25). However, when we looked closely at the horizontal sections of 1.5 μm thickness, we realized that cells were not distributed uniformly but there were many holes without cells (Figure 2B, z-section and Video S2). To see how these holes are formed, we visualized the cell-cell adhesion by staining for N-cadherin. It revealed that the mesoderm cells were connected via N-cadherin mediated cell-cell adhesion and surrounded the holes, which led to formation of a meshwork structure within the tissue (Figures 2B, 2C and 2D, and Video S2). Thus, in contrast to previous reports (18) that suggested that the mesoderm is densely packed with migrating cells, we found a meshwork structure of collectively migrating mesoderm cells. The holes in the meshwork structure were surrounded by about 10 to 20 cells (Figure 2C). The transverse section of the embryo revealed that the holes extended in the z-direction between the epiblast and the endoderm layers (Figure S2A).

**Figure 2.**
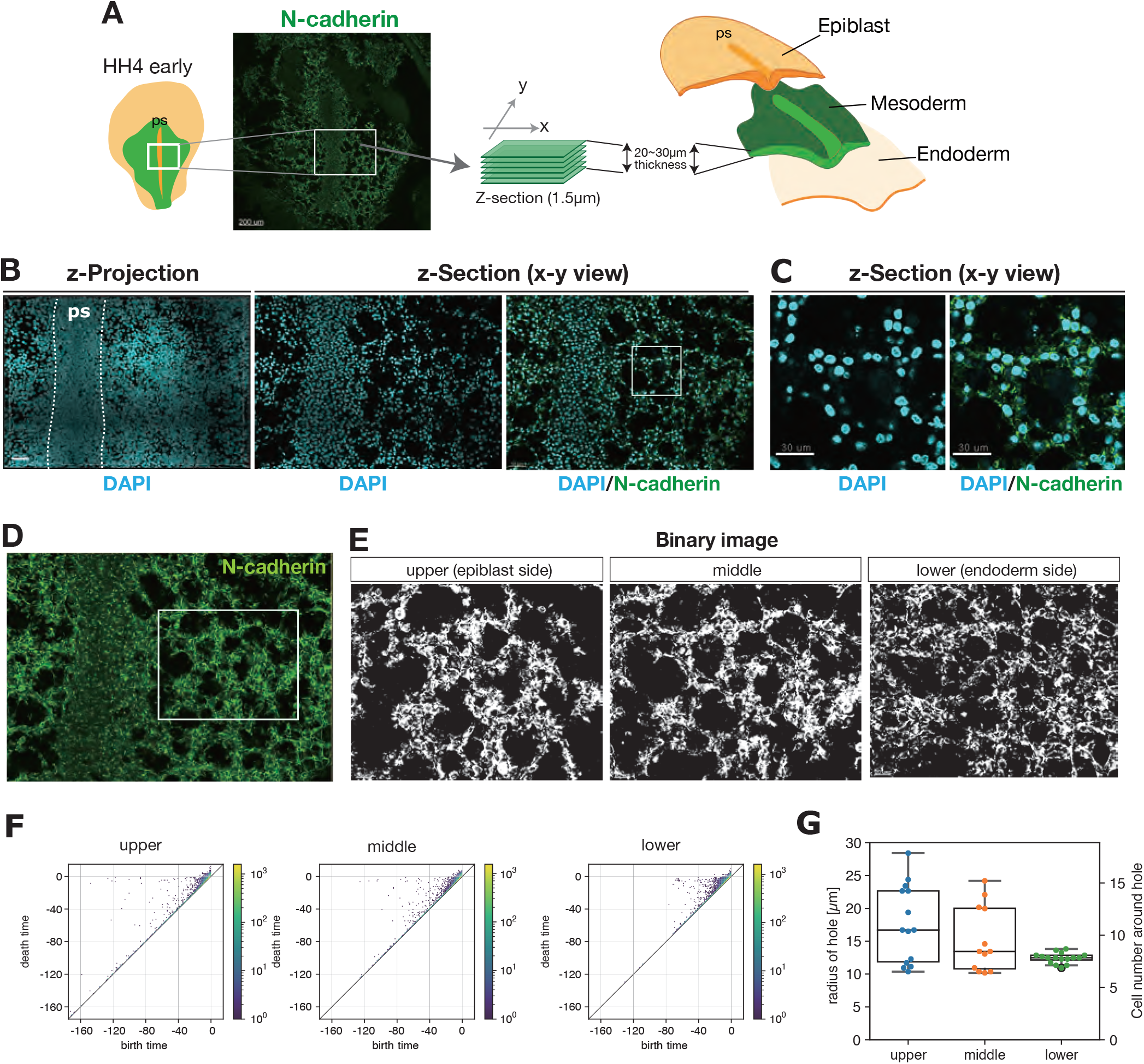
Meshwork structure in mesoderm during gastrulation. (A) Schematics of the 3D imaging. The white box indicates the imaged area shown in (B). (B) Spatial distribution of the cells in the fixed mesoderm tissue stained for nuclei (cyan) and N-cadherin (green) in the z-projection view (left) and the horizontal section (middle and right). (C) Magnified view of the characteristic meshwork structure in the white box in (B). (D) N-cadherin expression in the middle section of the mesoderm. (E) Binary images of three z-sections in the white box in (D). (F) Persistence diagram (PD) obtained by applying persistent homology analysis to the three z-sections in (E). The pixel size in (E) is 0.192 μm. The points forming a hole branch around death time ~ 0 correspond to the holes. (G) Statistics of the radius of holes that appear in the hole branch in the PD and the number of the cells surrounding the holes given that the cell diameter is 10 μm.

To characterize the meshwork structure of the mesoderm quantitatively, we applied persistent homology analysis (SI Methods). Persistent homology is a tool that has been developed recently in applied mathematics to quantitatively characterize the topological structure in disordered systems (26–28). In persistent homology analysis, the holes are characterized by the two quantities called birth and death times, and they are visualized by points in persistence diagram (PD) where the coordinates are given by the birth and death times (Figure 2F). The difference between the death time and the birth time is called lifetime, which becomes large for a reliable topological structure. Note that in the persistent homology on binary images, the birth and death times, and thus the lifetime, are measured in the unit of pixels. We performed persistent homology analysis by using a software named HomCloud (27) and applying it to binarized images of the mesoderm at three different z levels. The PDs in Figure 2F show the results correspond to the images in Figure 2E. The points with large lifetime were distributed around the death time ≈ 0 as a branch extended away from the diagonal line (Figures 2F and S2B), which we call *hole branch* in this paper. Each point in this hole branch corresponded to a clear hole in the binary image (Figure S2B). To characterize the size of the holes quantitatively, we calculated the radius from the birth time of a hole in the hole branch (Figure 2F, see also SI Methods), which was estimated on average as 17 μm, 15 μm, and 12 μm for the upper, middle and lower layers, respectively (Figure 2G). Thus, the diameter of hole is almost comparable to the characteristic length scale of collective migration measured in the previous subsection (~60 μm). To conclude, these results indicate that the mesoderm is extended as a loose layer of cells, and the characteristic meshwork structure is formed by cells connected by N-cadherin-mediated cell adhesion, which may lead to their collective migration.

### Meshwork structure is dynamic with the emergence and collapse of holes

To examine whether the meshwork structure was also observed in living embryos, we next performed live imaging of transgenic chicken embryos that ubiquitously express cytoplasmic GFP (29) to visualize the migration of the mesoderm cells and how they interact with each other (Figure 3A). We again found that cells away from the primitive streak formed meshwork structures, as seen in the optical thin section of middle layer in the mesoderm at the mid-streak level (Figures 3A, S3A, and Videos S3 and S4). Interestingly, the holes in the meshwork were not static but gradually moving anterior-laterally (Figures 3A, S3B, and S3C, and Videos S3 and S4). We also notice that a cell of one hole migrates and participates in another hole as time passes. (Figure 3A and Videos S3 and S4).

**Figure 3.**
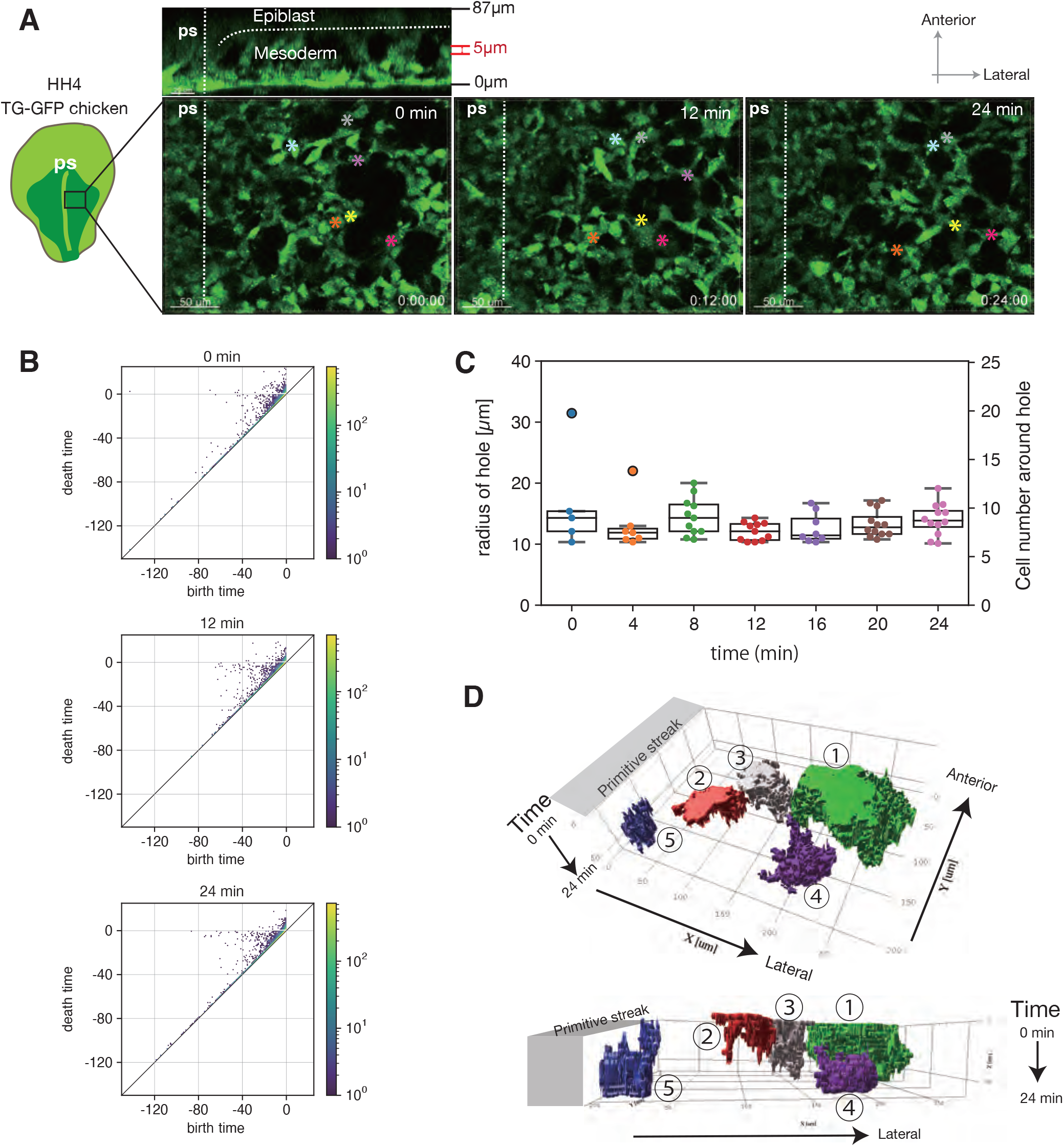
Dynamic meshwork structure. (A) Successive snapshots obtained from a live image of mesoderm tissue. The position of the 6 cells at different time points are indicated by the colored asterisk. (B) Persistence diagram (PD) of the three snapshots in (A). The hole branch of the points around death time ~ 0 away from the diagonal line. The pixel size in (A) is 0.22 μm. (C) The time series of the radius of holes that appear in the hole branch in the PD, and the corresponding number of the cells surrounding the holes that is calculated from the radius under the assumption that the cell diameter is 10 μm. The p-values between any two time points obtained by t-test were larger than 0.05 except for the pairs of 0 min and 8 min, 0 min and 12 min, 0 min and 16 min, 0 min and 20 min, 0 min and 24 min, 4 min and 12 min (p<0.01), and 8 min and 12 min (p<0.05), which might possibly be caused by the small size of the data set. (D) Spatiotemporal diagram of the holes. The holes were dynamic with the appearance (4) and disappearance (2,3) as well as the fusion (5) and fission (2).

To confirm this dynamic meshwork structure is the same structure as those found in the fixed embryos in the previous subsection, we applied persistent homology analysis to the snapshots of the time-lapse movie (Figure 3A). The PDs at different time points (0, 12, and 24 min) showed the hole branch of the points with large lifetime, which was comparable to the PDs obtained for the fixed embryo (Figures 2F and 3B). From this, we conclude that the meshwork structure found in the living embryo is the same structure as those found in the fixed embryos. The radius of the holes was on average about 13 μm, which showed no systematic change during the observation over 24 min. (Figure 3C).

Finally, we investigate how the meshwork structure changes in time. The advanced inverse analysis enables us to detect the region of the hole in the original binary image, which corresponds to each point in PD (SI Methods and Figure S3D). To visualize the time-evolution of the holes, we plotted the region of some holes obtained by the advanced inverse analysis in x-y-t coordinates over 24 min (Figure 3D). The holes underwent emergence and collapse, and they also split and merged occasionally, while moving gradually in the anterior-lateral direction (Figures 3D, S3B and S3C).

These results implied that the mesoderm cells are only transiently connected to each other and can easily change their partners. Thus, the meshwork structure is dynamic in the sense that cells that consists of the holes are replaced over time while the mesoderm cells migrate away from the primitive streak. Since the length scales of the polar order of the collective migration and the size of the holes of the meshwork are comparable, we speculate that the frequent rearrangement of the cell-cell contact is a reason why the collective order of the mesoderm cells decays in the long length scale (Figures 1I).

### Cell-cell adhesion is important for collective cell migration during mesoderm formation

Previous studies based on scanning and transmission electron microscopy observations have reported that the space between the epiblast and endoderm in early chick embryos, where the mesoderm cells migrate, is filled with water-soluble components such as hydrated glycosaminoglycans, in particular hyaluronic acid, and contains little extracellular matrix that can provide a scaffold for cell migration (30–32). This suggests that the mesoderm cells rely on cell-cell adhesion to get traction to migrate as well as the contact to the basal lamina of either epiblast or endoderm (31, 33). The formation of the meshwork and its dynamic characteristics may also rely on cell-cell adhesion. Taking these points into consideration, we questioned how much impact cell-cell adhesion has on cell migration as well as on the meshwork structure.

Consistent with the previous reports (12, 34, 35), during gastrulation, most mesoderm cells expressed the classical adhesion molecule N-cadherin (Figure 2A). Notably, we found that N-cadherin was localized at the cell-cell contact site between the cells in both the x-y and x-z sections (Figure 4A), meaning that the N-cadherin-mediated cell-cell adhesion is present at the cell-cell contact sites between upper and lower cells as well as at the horizontal cell-cell contact site. This implies that N-cadherin-mediated cell-cell adhesion plays fundamental role in the formation of the meshwork structure and the collective migration. To test this idea, we studied the effect of reducing the intercellular adhesion of mesoderm cells (Nieman et al., 1999; Ozawa, 2015; Ozawa and Kobayashi, 2014). To this end, we generated a deletion mutant of N-cadherin (N-cad-M) which lacks the extracellular (EC) domain that is responsible for adhesive activity (36–38). In addition, H2B-mCherry was flanked on its 3’-side of the 2A peptide to make the N-cad-M expressing cells detectable (Figure 4B). We over-expressed N-cad-M with H2B-mCherry in the mesoderm cells.

**Figure 4.**
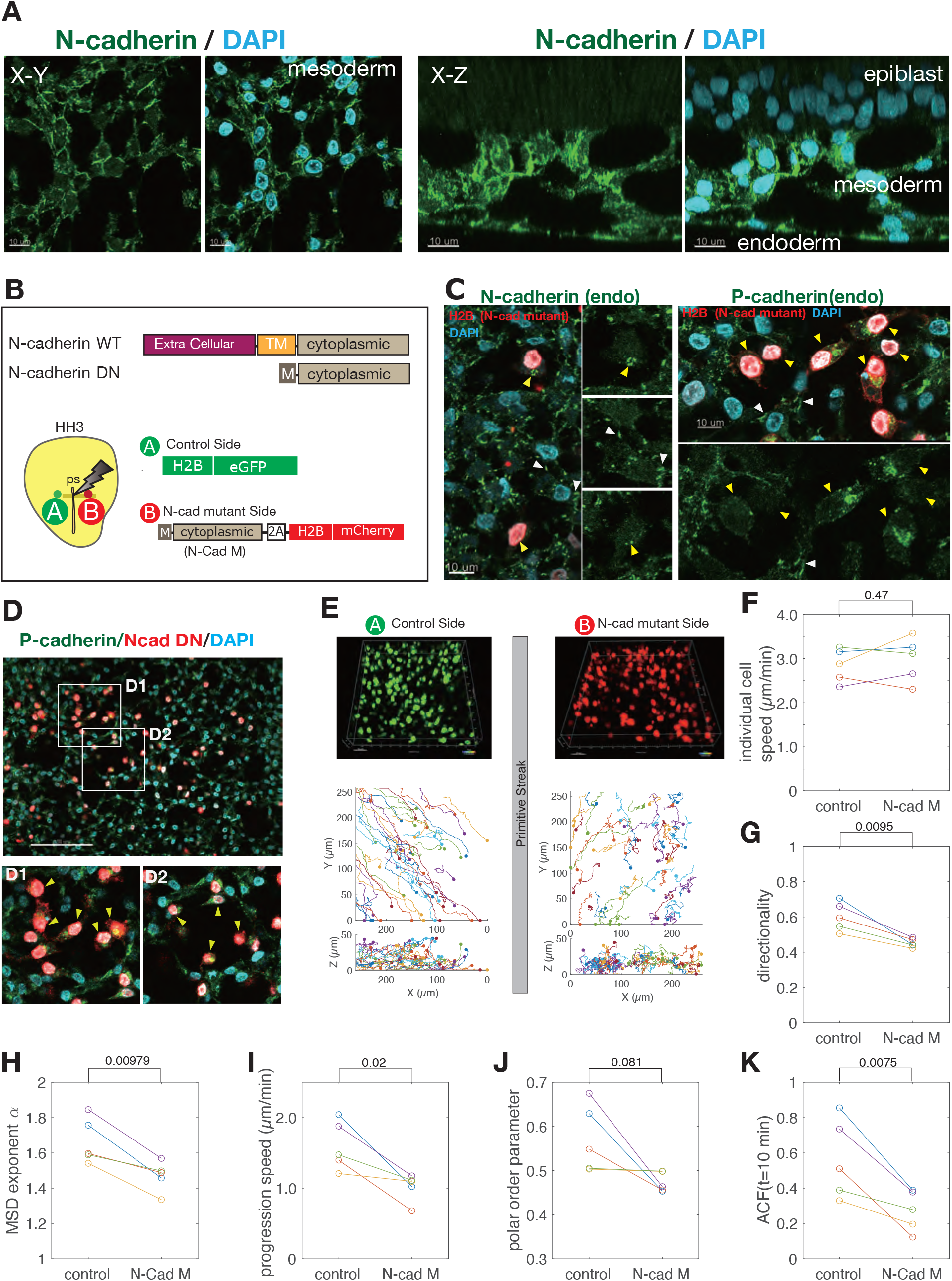
Intercellular adhesion controlling collective mesoderm cell migration. (A) N-cadherin expression in the mesoderm. N-cadherin was localized at the cell-cell contact sites both in the horizontal section (left) and in the vertical section (right) that surround the holes. Scale bars, 10 μm. (B) Structure of the wild-type N-cadherin and the deletion mutant of N-cadherin consisting of the cytoplasmic domain with myristoylation signal (top). Schematic diagram of the experimental method (bottom). To compare the migration of the mesoderm cells, H2B-eGFP was electroporated on the A side, while the N-cadherin mutant (N-Cad-M) was electroporated on the B side. The N-Cad-M expressing cells were marked by the H2B-mCherry expression. (C) Effects of N-Cad-M overexpression on endogenous cadherin expression. Endogenous N-cadherin (left) and P-cadherin (right) are expressed specifically at the cell-cell contact site in the control mesoderm cells (white arrow heads). In contrast, in the cells expressing N-Cad-M labeled in red, the expression of N-cadherin (left) and P-cadherin (right) were almost disappeared from the cell membrane (yellow arrow heads). Scale bars, 10μm. (D) N-Cad-M expressing cells are excluded from the meshwork structure of control mesoderm cells (yellow arrow heads). (D1) and (D2) Magnified images in the white boxes in the top panel. The N-Cad-M expressing cells did not participate in the meshwork. Scale bar, 100 μm. (E) Examples of (top) a snapshot of the live imaging and (bottom) trajectories of the mesoderm cells expressing H2B-eGFP (A side) and the N-cad-M (B side) of the same embryo. The initial position of the cells is marked by dots on the trajectories. (F-K) Statistical quantification of the migration behavior of the control and N-cad-M cells for five embryos. The corresponding statistical quantity of each cell in each embryo is shown in Figure S4. The quantities of control and N-Cad-M in the same embryo are linked by the line. (F) Mean of individual cell speed (*p*=0.47). (G) Mean of directionality (*p*=0.0095). (H) MSD exponent (*p*=0.00979). (I) Mean of progression speed (*p*=0.02). (J) Polar order parameter (*p*=0.081). (K) Auto-correlation function (ACF) of the direction of collective migration at 10 min (*p*=0.0075). *p*-values were obtained using paired t-test of the five embryos. The xy size of the imaged square area of 5 embryos: 258, 192, 207, 500, 500 μm. The numbers of cells analyzed: N=118 (Control), 119 (Mutant) (embryo 1), 41(C), 28(M) (embryo 2), 44 (C), 30 (M) (embryo 3), 253 (C), 290 (M) (embryo 4), 297 (C), 232 (M) (embryo 5).

To ensure that the endogenous N-cadherin was disappeared from the membrane, we used an N-cadherin antibody reactive against the EC domain, which can detect only endogenous N-cadherin. Immunostaining with this antibody showed that the endogenous N-cadherin accumulated in the cytoplasm and its expression was disappeared from the membrane of the mutant mesoderm cells (Figure 4C left, cells with nuclei labelled in red). Similarly, endogenous expression of P-cadherin was affected, which was also mainly detected in the cytoplasm of mutant cells (Figure 4C right, cells with nuclei labelled in red). From these results, we suppose that the preferential localization of large amounts of N-cad-M to the plasma membrane disrupted the membrane localization of endogenous cadherins, which effectively attenuates cadherin-mediated cell-cell adhesion. Indeed, the cells expressing N-cad-M were excluded from the meshwork of the control cells (Figure 4D). In addition, the mutant cells were less elongated and exhibited more rounded shape (Figures 4C and 4D).

Using this mutant form of N-cadherin, we performed live imaging to investigate the effect of the cell-cell adhesion on the mesoderm cell migration. To compare with the control cells, we electroporated the N-cad-M construct into the mesoderm cells on one side of the primitive streak, and we introduced a plasmid expressing H2B-eGFP to the cells on the other side to trace them as control cells (Figures 4B and 4E, Video S5). We performed this analysis for five embryos, each of which contains a lot of cells (see Figure 4E-K and Figure S4). We found that the cells expressing N-cadherin mutant exhibited meandrous motion with more frequent changes in migration direction than the control cells (Figure 4E, Video S5), although the migration speed along the trajectory was not statistically different between them (Figures 4F and S4A). This is confirmed by the significant reduction of the directionality (Figures 4G and S4B) and the smaller exponent of the MSD for the mutant cells (Figure 4H). In addition, the progression speed of mutant cells was lower than that of the control cells for each embryo (N=5) (Figure 4I and S4C), indicating that the progression of mesoderm tissue became slower when the cell-cell adhesion was attenuated. These results show that the cell-cell adhesion mediated by N-cadherin plays an important role in the tissue progression and the directionality of the mesoderm cell migration.

We next investigate the impact of cell-cell adhesion on the collective migration of the mesoderm cells. The polar order parameter *φ*(*t*) on the mutant side was smaller than that on the control side in three embryos out of five (Figure 4J and S4D), suggesting that the alignment of the cell migration direction is controlled by cell-cell adhesion. To see the persistence of the direction of collective migration, we measured its auto-correlation function (ACF) (SI Methods). The ACF of the direction of collective migration for the mutant cells decayed faster than that of the control cells in all five embryos (Figures 4K and S4E), indicating that the mutant cells changed the direction of collective migration more frequently than the control cell did. These results indicate that the cell-cell adhesion maintains the collective migration of the mesoderm cells and the persistence of its direction.

To understand the difference in the properties of collective migration of the mesoderm cells, we carefully observed the time-evolution of the intercellular contact. The time lapse images showed that the control cells elongate their cell bodies and contact with each other via protrusions (Figure S5A). These cells were in contact for more than a few tens of minutes, and the longest contact duration lasted for more than one hour (No.1 and No.2 pair in Figure S5A upper panels, Video 6). In contrast, the mutant cells did not maintain their cell-cell contacts upon collision with other cells for more than 20 minutes. (See No.1 cell in Figure S5A bottom panels, Video 6). Thus, we speculate that the decrease in the contact time of the mutant mesoderm cells makes their motion random and the direction of the collective migration change frequently.

Taken together, the comparison of the migration characteristics between the control and N-cadherin mutant cells showed that the intercellular adhesion promotes the directionality of the individual mesoderm cells, their collective migration, and the tissue progression speed of the mesoderm (Figure 4F–K). We also found that the mutant cells were more rounded than the control cells and were excluded from the meshwork structure formed by the control cells (Figure 4D). We therefore hypothesize that cell-cell adhesion is one of the key factors for the formation of the meshwork structure. Unfortunately, for technical reasons, it was difficult to introduce N-cad-M into all mesoderm cells to see if tissues composed only of N-cad-M cells fail to form a meshwork structure. Therefore, we next tested our hypothesis by developing an agent-based theoretical model.

### Theoretical model to investigate the formation of meshwork structure

From the experiment using the mutant form of N-cadherin, we hypothesized that the cell-cell adhesion is one of the key factors for the meshwork structure formation. To understand how the dynamic meshwork structure of mesoderm cells emerges and how it is influenced by the cellular and intercellular properties, we develop an agent-based theoretical model. To this end, we modeled a cell by a rod-shape particle that interacts with others by short-range attraction with a repulsive core (SI Methods). To focus on the essential aspect of the meshwork formation without complication, we will consider a model in two-dimensional space. We note that previous theoretical studies reported that elongated cells with attractive interaction can reproduce the formation of angiogenetic network structure (39, 40).

To start with, we studied the steady-state spatial distribution of agents, starting from the initial state where the agents were randomly positioned in space with random orientation. We investigate the impact of the cell-cell adhesion by changing the strength of the attractive interaction. To highlight the role of the attractive interaction, we kept the aspect ratio *γ* = 2 and omitted the self-propulsion of the agents. When the attractive interaction was absent or small, the agents were distributed randomly without any clear spatial pattern (Figure 5A, *ϵ*_atr_ = 0,0.001). However, for a sufficiently large attractive interaction, the agents formed a meshwork structure (Figure 5A, *ϵ*_atr_ = 0.01,0.05). The birth time calculate by the persistent homology analysis, which corresponds to the radius of holes, increased with the strength of attractive interaction (Figure 5B). These results demonstrate that the cell-cell adhesion plays an important role in the meshwork structure formation.

**Figure 5.**
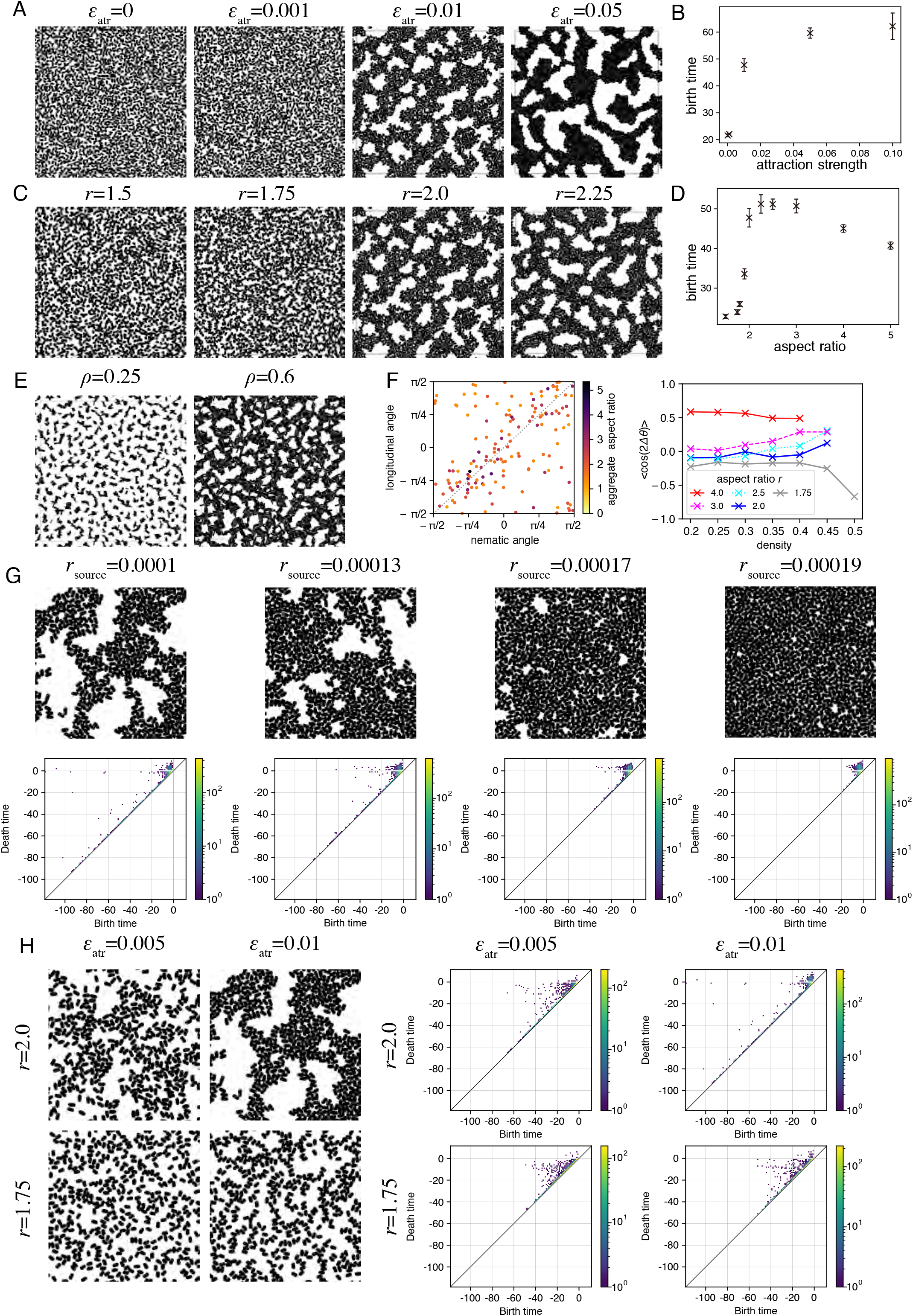
Theoretical model of meshwork formation. (A) Impact of the attractive interaction strength *ϵ*_atr_ on the meshwork structure formation. (B) Dependence of the birth time of the holes on the attractive interaction strength *ϵ*_atr_. The error bars indicate the standard error of mean obtained from n=10 independent simulations. (C) Impact of the agent aspect ratio r on the meshwork structure formation. (D) Dependence of the birth time of the holes on the aspect ratio *r* obtained from the persistent homology analysis. The error bars indicate the standard error of mean obtained from n=10 independent simulations. (E) Alignment of the agents in the aggregates and the meshwork structure. Aspect ratio *r*=4. (F) Relation between the nematic direction of agents in the aggregates and the elongation direction of the aggregates. (left) Relationship between the nematic angle and the longitudinal angle of each aggregate. The color of the points represents the aspect ratio of the aggregates. (right) The correlation between the nematic angle *θ_n_* and the aggregate longitudinal angle *θ_a_* defined by 〈cos2(*θ_n_* – *θ_a_*)〉 as a function of the aggregate density for different aspect ratio *r*. (G) Impact of the agent supply rate on the meshwork structure formation in the simulation with the agents supplied from the PS boundary on the left. Snapshots (top) and persistence diagrams (PD) (bottom). (H) Impact of the adhesion and the aspect ratio on the meshwork structure formation in the simulation with the agents supplied from the PS boundary. Snapshots (left) and PD (right).

Since the wild-type cells were more elongated than the mutant cells, we next studied the impact of the aspect ratio for sufficiently large attractive interaction, i.e., *ϵ*_atr_ = 0.01. When the aspect ratio *r* was small, the agents were distributed randomly without any spatial pattern (Figure 5C *r* = 1.5,1.75). However, there was a threshold aspect ratio *r*\* (1.75 < *r*\* < 2), beyond which the agents formed a meshwork structure with many holes void of agents (Figure 5C *γ* = 2,2.25). The persistent homology analysis distinguished this difference; at the threshold aspect ratio *r*\* ≈ 1.9, the birth time of the holes, which corresponds to the radius of hole, showed a sharp increase (Figure 5D). This transition is evidence that the elongated shape with a large aspect ratio is important for the meshwork structure formation.

In summary, our *in silico* results confirmed that both attractive interaction and elongated shape with a large aspect ratio are necessary for the formation of meshwork structure. Indeed, the aspect ratio of mesoderm cells was 2.34 ± 0.08 (± SEM) (Figure S5B, control), while that of the N-cadherin mutant cells was 1.91 ± 0.08 (± SEM) (Figure S5B, N-Cad-M). We emphasize that this experimental result is consistent with the existence of the threshold aspect ratio at *r*\* ≈ 1.9 in our *in silico* result (Figure 5D).

### Mechanism of meshwork formation

To understand the mechanism of the formation of meshwork structure, we focused on the small aggregates of agents that were found when the agent density was low. Since the results were quantitatively clearer for a high aspect ratio, we first set the agent aspect ratio as *r* = 4. When the density of agent is low, many small aggregates are formed due to the short-range attractive interaction (Figure 5E *ρ* = 0.25). Most of the aggregates have an elongated shape with aspect ratio much greater than unity (Figure 5F left, color). Inside the aggregates, the agents tend to align their direction of the shape elongation. Such a directional order is called nematic order. The direction of the nematic order in each aggregate is correlated with that of the aggregate elongation (Figure 5F left and the red curve in Figure 5F right), indicating that the aggregates tend to elongate in the direction of the nematic order. When the agent density increases beyond a threshold value, a meshwork structure is formed (figure 5E *ρ* = 0.6). Thus, a scenario for the formation of the meshwork structure is as follows. The attractive interaction between agents induces the formation of aggregates, in which the agents align their orientation nematically. The positional and directional fluctuations of the agents deform the shape of aggregates in a way that they elongate in the direction of the nematic order. As the agent density increases, such elongated aggregates further extend and are eventually connected to each other, leading to the formation of the meshwork structure due to the randomness of the aggregate elongation direction. When the aspect ratio *r* is reduced to *r* = 2, the correlation of the orientations of the nematic order and the aggregate elongation decreases (the blue curve in Figure 5F right). As the agent density increases, however, the correlation increases, indicating that the same scenario applies to this case (*r* = 2). In contrast, when *r* = 1.75 < *r*\*, the correlation decreases as agent density increases, suggesting that the scenario does not hold, resulting in the random distribution of agents without the meshwork structure formation.

### Dynamic meshwork formation with the supply of agents

During gastrulation, mesoderm cells are continuously supplied and move away from the primitive streak (Figure 1A). To mimic this situation, we modified the simulation condition to the case where the agents are supplied constantly at the rate from one side of the boundary, which corresponds to the primitive streak (SI Methods). We also switched on the self-propulsion of the agents in the direction of the shape elongation and introduced a slight chemotaxis so that the agents move efficiently away from the primitive streak boundary (PS boundary). When the supply rate was high, the space was filled by the agents leaving no clear holes (Figure 5G, *r_source_* = 0.00019). In contrast, when the rate was decreased, small holes appeared, the size of which increased as the supply rate was further decreased (Figure 5G). These holes move away from the PS boundary in the lateral direction. The holes also showed dynamic behaviors, such as emergence, collapse, splitting and merging, due to the agent self-propulsion (Figure 5G, *r_source_* = 0.0001, and Video S7), which resembled the experimental observations of TG-chick embryos (Figure 3A, S3B, and S3C, and Video S3 and S4). To characterize the structure quantitatively, we performed persistent homology analysis and found that the points of birth-death pairs with large lifetime were distributed in a hole branch around the death time ~ 0 when the supply rate was small (*r_source_* = 0.0001, Figure 5G, bottom), consistent with the experimental observation (Figure 2F and 3B). As the supply rate increases, the holes void of cells became smaller and, correspondingly, the hole branch shrank. In consequence, when the supply rate *r_source_* = 0.00019, the points were distributed in a clumped pattern near the diagonal line (Figure 5G, bottom). These results indicate that the supply rate, which controls the density of the cells in the mesoderm, is an additional important parameter for the meshwork formation. Note that the agents cannot form a meshwork structure at a very low density (Figure 5E). We also confirmed that the decrease in either the aspect ratio of agent shape or the attractive interaction prevented the formation of the meshwork structure when supply rate was *r_source_* = 0.0001 (Figure 5H).

### Dependence of mesoderm meshwork structure on the developmental stage

Now a new question arises: How do the meshwork structures of the mesoderm cells change during the embryonic development? To answer this question, we performed a persistent homology analysis using the horizontal slice images at different developmental stages. Interestingly, we found that as the developmental stage proceeds, the size of holes decreases and eventually the space was filled by cells (Figure 6A, top). The corresponding PDs showed the points of birth-death pair with large lifetime were distributed in a hole branch around death time ~ 0 at HH3+. However, as the developmental stage proceeded to HH4+, the hole branch shrank, and the distribution of the points was eventually changed to a clumped pattern near the diagonal line (Figure 6A, bottom). The average radius of holes was reduced from about 8 μm at HH3+ to 5 μm at HH4+, which took about 6 hours (Figure 6C). Note that the average radius of the holes at HH3+ becomes 15 μm when we focused on the larger holes by setting the threshold birth time to the same as that for Figures 2G and 3C (see also SI Methods). Thus, while the size of holes is maintained for about half an hour (Figure 3C), it gradually decreases over several hours. From the simulation results shown in Figure 5G, we speculated that the supply rate of mesoderm cell from the primitive streak increases gradually as the developmental stage proceeded. Therefore, we performed a simulation with a time-dependent supply rate of the agent from the PS boundary (SI Methods). We found that the size of the holes was initially large, but gradually decreased with time (Figure 6B). The radius of holes obtained from the PDs (Figure 6B bottom) decreases as well (Figure 6D), although the hole sizes in the simulation were slightly larger than those of the experiment. This quantitative difference might come from the fact that the shape of the cells in the simulation is kept constant, while the shape of the real mesoderm cells changes dynamically. Nevertheless, from these results, we conclude that one possible reason of the decrease of the hole size as the developmental stage of the embryo proceeds is the increase in the rate of the appearance of the mesoderm cells at the primitive streak.

**Figure 6.**
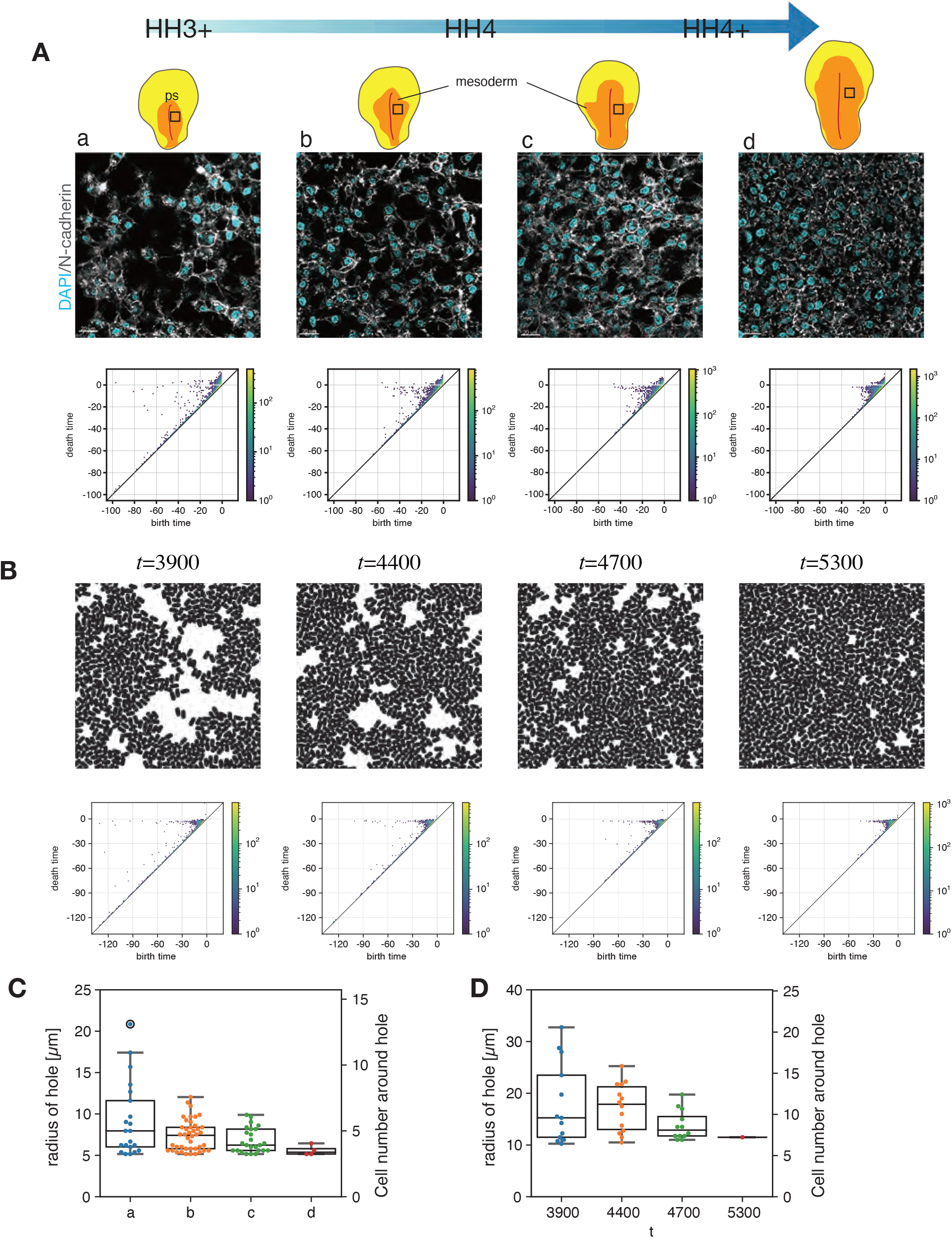
Changes in the meshwork structure during development. (A) Spatial distribution of the cells in the mesoderm tissue stained for nuclei (cyan) and N-cadherin (white) at different developmental stages (top) and the corresponding persistence diagrams (PD) (bottom). The persistent homology analysis was performed using binary images. The pixel size is 0.215 μm. (B) Simulation with the supply of the agents where the supply rate increases with time. Snapshots (top) and the corresponding PD. (C) The radius of holes that appear in the hole branch in the PD in (A). (D) The radius of holes that appear in the hole branch in the PD in (B).

## Discussion

Our results revealed a novel mode of collective cell migration, in which the migrating nascent mesoderm cells form a dynamic meshwork structure in three-dimensional space between the epiblast and endoderm while moving collectively in the anterior-lateral direction. In the early gastrulation stage of chick embryos, the fate of the various cell populations in the mesoderm has been studied in detail and it is known to be determined by the final migration destination (15). However, it was not well understood whether the mesoderm cells move collectively without scattering toward their destination. In addition, the mesoderm cells were thought to be densely packed without any spatial structure (15, 19). In this study, we investigated these points quantitatively by the 3D time lapse imaging and the horizontal thin optical sectioning of the mesoderm in fixed whole mount embryos applying tissue clearing method. From the analysis of the multicellular tracking data, we confirmed that the mesoderm cells migrate collectively with the characteristic decay length of about 60 μm. In addition, from the horizontal thin sections, we found that the mesoderm cells form a meshwork structure. The diameter of the holes is about 30 μm, which is almost comparable to the characteristic decay length of the collective migration. From these results, we presume that this meshwork structure is relevant to the collective migration of the mesoderm cells. Since only little extracellular matrix exists in the mesoderm (30–32), the formation of meshwork structure should be based on the intercellular adhesion. In fact, the disruption of the intercellular adhesion using a mutant form of N-cadherin resulted in the exclusion of the mutant cells from the meshwork of the control cells. Moreover, although the migration speed along trajectory is unaltered, the directionality of individual cell migration, the tissue progression speed, and the stability of the direction of collective motion were reduced for the mutant cells compared to the control cells. These results indicate that the cell-cell adhesion coordinates the migration of the mesoderm cells. To summarize, we conclude that the cell-cell adhesion plays a fundamental role in the meshwork formation for the mesoderm cells to migrate collectively. Such collective motion could contribute to the robust formation of cell migration pattern in response to guidance signals such as chemoattractant and chemorepellent (19).

Extracting information about the organization and arrangement of cells in tissues from microscopy images and comparing them with mutants has been largely based on visual inspection. Moreover, their quantitative and objective characterization is often challenging because of their variability and lack of periodicity. To obtain the information of the holes of the meshwork structure in the mesoderm tissues such as their size and position objectively and automatically, we used persistent homology, a tool of topological data analysis (TDA). TDA is a recently growing unique methodology, and it provides geometric information of the complex data, which has been employed in physical, medical and biological research (28, 41–43). We used this method to extract the information of the dynamics of the meshwork structure, which is still a challenging task in TDA. By applying the same analysis method to the simulation result, we compared it with the experiment quantitatively and we successfully showed that the theoretical model captures the essential aspect of the meshwork formation observed experimentally.

The *in vivo* collective migrations of mesenchymal cell have been reported for the neural crest cells in frog and chick (8, 44). These neural crest cells migrate on a two-dimensional surface within a confined space with a physical barrier of neighboring tissues (8). Contact attraction (44) and contact inhibition (45) orient the cell motion to induce the collective cell migration such as chain migration and stream formation (46). In contrast, mesoderm cells at the early gastrulation stage migrate in the three-dimensional space between the epiblast and endoderm without physical barrier in the lateral direction. Almost all cells were attached to other cells. Upon collision, the mesoderm cells stay in contact for more than a few tens of minutes (Figure S5A). Thus, contrary to the neural crest cells, the mesoderm cells did not show contact inhibition of locomotion, which is consistent with the case of the mouse mesoderm cells (47). In situations where cells exploit other cells as scaffolds and the cell density is low, we speculate that forming a meshwork rather than a three-dimensional mass would be more efficient to extend the distance.

In the nascent mesoderm tissue of chick embryo, matured ECM is almost absent in the intermediate layer where cells are in contact with other cells but not with the epiblast or endoderm (30–32). How cells in the intermediate layer can generate traction force for the movement is an intriguing future question. Mesoderm cells on the basal lamina of either epiblast or endoderm can generate traction force to migrate. By adhering to these cells, it may be possible that the mesoderm cells in the intermediate layer move forward together in a passive manner. In addition to such passive movement, the intermediate cells might generate active force at the intercellular contacts, by which they migrate further. Another possibility for the active process of the cell motility is the treadmilling of intercellular junction, which has been implicated in the migration of adhering cells (48).

During the vasculogenesis, endothelial cells also form a meshwork structure with cords of cells that surround the regions void of cells. In this case, cell aggregates formed initially are connected to organize into a primitive vascular plexus (49). The cell motions appear to be random along the cords (50). In contrast, the meshwork structure we observed is formed by the mesoderm cells which are provided from the primitive streak without the formation of cords of cells and a different lineage than the cells that contribute to the vasculogenesis. Moreover, the mesoderm cell motion is biased to the anterior-lateral direction. Thus, although there are some similarities in the 2D horizontal section patterns, the 3D structures are different between these two cases.

During the development of enteric nervous system, enteric neural crest cells (ENCCs) migrating in the mesenchyme also form a meshwork structure within a narrow 2-dimensional layer (51). ENCCs migrate in chains and the cells immediately behind the preceding chains often follow the same path (52). Thus, the network created by the preceding cells often remained intact for many hours. This constant shape of the network contrasts with the dynamic properties of the meshwork structure formed by the mesodermal cells in the chick embryo.

To understand how the meshwork forms, we developed a theoretical model that demonstrated that the elongated shape of agents and the attractive interaction between them are the key factors for the formation of meshwork. While a previous study reported that the branches of a meshwork structure showed nematic order (39), it was not clear how a meshwork structure emerges as the density increases. We showed quantitatively that clusters composed of agents deforms in the direction of the nematic order of the agent elongation. As the density increases, the elongated clusters grow and finally fuse with each other to form a meshwork structure. Although the agents in the current model do not deform their shape, actual mesoderm cells do, which enables them to migrate. Presumably, the shape deformation may play an important role for the 3D meshwork structure formation where the intermediate cells have no scaffold to migrate other than other cells, like the one that we found in the mesoderm of chick embryo. It is thus a future work to investigate how the shape deformation of the agents contributes to the 3D meshwork structure formation.

## Acknowledgements

We thank Guojun Sheng for critical reading of our manuscript and the members of Laboratory for Physical Biology for discussions. This work was supported by Kakenhi grant 16K07385 (YN), 19H14673 (MT) and 22H05170 (TS), JST CREST Grant JPMJCR1852 (TS) and the core funding at RIKEN Center for Biosystems Dynamics Research (TS).

## Materials and Methods

### Chick Embryo Collection and ex vivo culture

Fertilized hen’s egg (Shimojima farm, Kanagawa, Japan) or fertilized transgenic chicken’s eggs (Avian Bioscience Research Center at Nagoya University) (Table S1) were incubated at 38.5°C until embryos reached the desired developmental Hamburger-Hamilton stage (20).

### Electroporation

For electroporation, expression vectors were injected between the epiblast and vitelline membrane of embryos at a concentration of 2-5ug/ul and electroporated with 1mm platinum electrodes by using an electroporator (NEPA21 Super Electroporator; Nepagene) with the following parameters: 8.0 V, 0.5 ms width, one poring pulse, followed by 5.0 V, 25.0 ms width, 50 ms interval, five polarity exchanged transfer pulses. Embryos were then cultured for several hours according to the Easy Culture (EC) protocol (21).

### Generation of chick-N-Cadherin mutant

Full-length of N-cadherin coding sequence (accession number NM_001001615.1) was amplified by PCR from cDNA of HH 5-7 chick embryos using the following primers: Fw 5’-ATGTGCCGGATAGCGGGAAC-3’ and Rev 5’-TCAGTCATCACCTCCACCG-3’, which was subcloned into the pGEM-T Easy vector (Promega). The full-length of N-cadherin fragment was then used as a template to generate an N-cadherin mutant lacking the extracellular and transmembrane domains, which corresponds to amino acids 752-912 of the N-cadherin protein. To ensure membrane localization of N-cadherin mutants, the second PCR reaction was performed using following primers in which the sequence of an N-myristoylation signal from Src kinase (53) was added at the 5’ side of the forward primer: Fw 5’-ATGGGTTCTTCTAAATCTAAACCAAAAGATCCATCTCAACGTATGAAGCGCCGTGATAAGG-3’, and Rev 5’-GTCATCACCTCCACCGTAC-3’. This amplified fragment was named N-Cad-M and was subcloned into the 5’ side from P2A peptide (ATNFSLLKQAGDVEENPGP) of the pCAG-P2A-H2B-mCherry vector by In-Fusion Cloning (Takara, Japan). To visualize the membrane of cells that express N-Cad-M, the N-Cad-M-P2A was amplified by PCR with a DNA fragment set of 5’-GCGGCCGCGGATCCGCATGCGCCACCATGGGTTCTTCT-3’ and 5’-TTGCTCACCATAACGCATGCTTTAGGTCCAGGGTTCTCC-3’, which was then subcloned into the 5’ side from the eGFP sequence of the pCAG-eGFP-CAAX-P2A-H2B-mCherry expression vector by In-Fusion Cloning. The above oligonucleotides used in this study are listed in Table S2 and the recombinant DNA constructed in this study are summarized in Table S3.

### Immunohistochemistry

For immunohistochemistry, embryos are fixed in 4%PFA, and the following antibodies were used: Purified Mouse Anti-E-Cadherin (610181, BD transduction Lab); Anti-N-cadherin, polyclonal (Code No. M142, Takara bio); Monoclonal Anti-N-Cadherin/A-CAM (Clone GC-4, Product No. C2542, Sigma-Aldrich). Alexa Fluor secondary antibody (Goat anti-Rabbit IgG, Alexa Fluor 488, A-11034; Goat anti-Mouse IgG, Alexa Fluor 488, A-11029, Thermo Fisher Scientific) were used for double color detection. DAPI (Cellstain DAPI Solution, 1:100, 340-07901 Dojindo Laboratories) for the labeling the nucleus was used. After washing, embryos were cleared with SeeDB-2G solution (54) before being processed for imaging. Immunofluorescence images were captures with a laser scanning confocal microscope (FV3000RS with IX83 inverted; Olympus) equipped with UPLSAPO 30xS/1.05 NA, 60xS/1.3 NA objective lenses, using Fluoview (Olympus) as the image acquisition software. For each embryo, several images corresponding to different focal planes and different fields were captured using z-section and tilling functions. The acquired images were imported to Imaris 9.5.1 (Oxford instruments, UK) to 3D-visualize for further analysis. The antibodies used in this study are summarized in Table S4.

### *in vivo* live imaging

For *in vivo* live imaging, the H2B-eGFP-expressing WT chick embryos or the transgenic-GFP chick embryos was transferred dorsal side up on glass-base dish (Iwaki, 3910-035) with semi-solid albumin/agarose (0.1%). Embryos were imaged at 38.5°C using an inverted multi-photon microscopy (Olympus MP, FVRS-F2SJ) coupled to a Maitai DeepSee HP laser at 890 nm weave length and InSight DeepSee laser at 1100nm using 25x/water 1.05 NA long distance objective lens (XLPLN25XW-MP).

### Obtaining the trajectory of individual mesoderm cells

To obtain the trajectories of mesoderm cells, live imaging data of embryo expressing H2B-eGFP were analyzed using IMARIS (Oxford Instruments). The movement of each nucleus was identified using *“Spot”* function in the package “IMARIS for tacking” as described below. For identifying the nuclear position, we used *“Spot Detection”* with the parameter *“Estimated Diameter”* to be 6 μm by adjusting the lowest threshold in *“Quality”* setting in the *“Filter”* section to a value with which the faintest nuclei were reliably distinguished from the background. For tracking, *“Autoregressive Motion”* were used in the *“Algorithm”* section with *“Max Distance ”* to be 8μm and *“Max Gap Size ”* to be 3 without using *‘Fill gaps with all detected objects’*. We then removed the short tracks by applying the *“Track Duration* above 1800 s” in the *“Classify Tracks”* section. The trajectory data obtained in the above way was then exported as a comma-separated values (csv) file for the further analysis. Then, the mean square displacement, directionality, polar order parameter and mean square relative distance as described in the following sections were obtained using a custom-made code of Matlab (Mathworks Inc., Natick, MA).

### Individual cell speed

The instantaneous velocity of each cell is defined by the displacement of the cell position in two subsequent images divided by the time interval. The individual cell velocity is calculated by averaging the instantaneous velocity over the trajectory. The individual cell speed is the magnitude of the individual cell velocity.

### Progression velocity and progression speed

To calculate the tissue progression speed, the image window is divided into small regions of 50 μm x 50 μm as in Figure 1D. Then, the progression velocity is calculated as the temporal average of the average velocity of cells in each region at each time point. The progression speed is the magnitude of the progression velocity.

### Directionality

The directionality was calculated using the formula given by

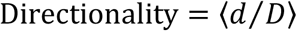

where *d* is the start-to-end distance and *D* is the actual length of trajectory between the start point and the end point. The bracket 〈·〉 indicates the average over the trajectories of the cells in a sample. The value of directionality depends on the time interval of the trajectory. In this paper, we consider the trajectories for 20 min. The directionality is close to unity when the motion is in a straight trajectory, while it is close to zero when the motion is random or when the trajectory forms a closed loop.

### Mean squared displacement

For each sample, the mean squared displacement (MSD) was calculated for individual migrating cells and then average them over ensemble. The MSD for a given sample was calculated using the formula, given by

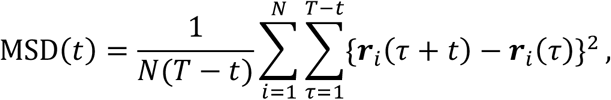

where ***r**_i_*(*τ*), *t, T*, and *N* are the 3D position of cell *i* at time τ, the lag time, final time, and number of trajectories in the sample, respectively. To obtain the exponent *α* of MSD, we fitted MSD(t) with the curve *Dt^α^* where *D* is a coefficient. For the fitting, we used lsqcurvefit of Matlab R2021b (mathworks). The exponent *α* is 1 for random motion, while it is close to 2 if the motion is ballistic (straight). For 1 < *α* < 2, the motion is known as super-diffusion.

### Auto-correlation function of velocity

For each sample, the auto-correlation function (ACF) of velocity was calculated for individual migrating cells and then average them over ensemble. The ACF of velocity for a given sample was calculated using the formula, given by

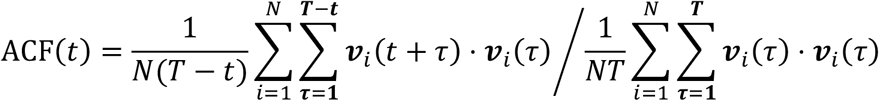

where ***ν**_i_*(*τ*), *t*, *T*, and *N* are the velocity vector of cell *i* at time τ, the lag time, final time, and number of trajectories in the sample, respectively. The ACF of velocity approaches to zero for sufficiently long time if there is no bias in the migration direction.

### Polar order parameter

For the trajectories obtained by the tracking analysis, the polar order parameter at a given time was calculated using the formula, given by

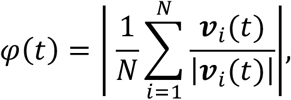

where *N* is the number of tracked cells, and ***ν**_i_*(*t*) is the instantaneous cell velocity of cell *i*. The polar order parameter *φ*(*t*) is close to unity if all cells move in the same direction, while it is close to zero if cells move in a random direction. For the data shown in Figure 1J, we calculated the temporal average of *φ*(*t*) in entire region (500μm x 500μm x z-depth). For the data in Figures 4I and S4C, since the size of imaged region was different between embryo samples, we divided the imaged region into subareas (125μm x 125μm x z-depth), in each of which we measured the temporal average of *φ*(*t*). Then, they were averaged in each embryo.

### Mean squared relative distance (MSRD)

We took a pair of cells which were initially at the distance less than 20μm, supposing that these cells were in contact with each other at that moment. Then, the mean squared relative distance (MSRD) was calculated for the pairs using the following formula,

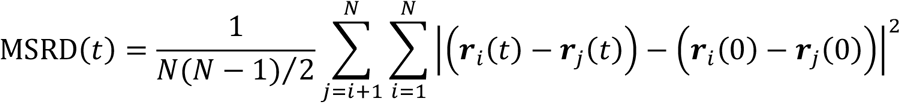

where ***r**_i_*(*t*), and *N* are the 3D position of cell *i* at time *t*, and number of trajectories in the sample, respectively.

### Topological structure analysis using persistent homology

To characterize the meshwork structure in the mesoderm quantitatively, we focused on the holes void of cells. To this end, we performed persistent homology analysis by using the software named HomCloud (3.0.1) (27). We first prepared a black and white binary pixel image from the original image by thresholding, which was used as an input data for HomCloud. Each topological structure is characterized by a pair of two values called birth and death times based on Manhattan distance, and thus, they are given in the unit of pixel. These two quantities basically represent the size of the identified topological structures and the distance between two topological structures, respectively. For the detailed explanation of the concept of the birth and death times, see (27). In HomCloud, the identified topological structures are visualized in persistence diagram (PD), which plots each pair of birth and death times. Since holes are identified by the 0th persistent homology, we focused on 0th PDs, i.e., the PDs for the 0th persistent homology. Each birth-death pair characterizes a black region in the binary image (Figure 2E)(Obayashi et al., 2018). The difference between the death and birth times is called lifetime. The topological structures with small lifetime are basically noise (27). Although in the original PD the points correspond to actual holes appear in the region with birth time < 0 and death time > 0, the death time of most holes in the experimental image becomes slightly smaller than 0 due to fluctuations possibly caused by several factors including those in staining and fluorescent imaging. By taking this into consideration, we identified the points with the birth time smaller than −10 μm and the death time larger than −2.5 μm as detected holes, except for the those in Figure 6C where the threshold is set as the birth time smaller than −5 μm and the death time larger than −2.5 μm because there was no hole satisfying the above stricter threshold for the later stage (Figures 6Ac and 6Ad). Since the magnitude of birth time corresponds to the shortest distance from the center to the periphery of a hole, we regarded this multiplied by the length of a pixel as the radius of the hole. The number of cells that surround each single hole was calculated from the perimeter length by assuming that a cell diameter is 10 μm.

### Analyzing the dynamics of holes

To visualize the spatiotemporal dynamic of holes, we first carried out the inverse analysis by HomCloud (27) for a total of 13 images at 2 min intervals out of the live imaging of 24 min and saved them as a series of images (Figure S3D bottom). From this 2D image sequence, we constructed a z-stack image using the 3D image reconstruction function of IMARIS by setting the z-interval at 5μm. Each hole was visualized by the *“surface”* function in IMARIS. To ensure each hole was visualized individually and the adjacent holes were reliably split, we used the following parameters. We set *“Threshold”* to 130, enabling *“Split touching Objects (Region Growing)”* and the value of the *“Estimated Diameter”* to 10 μm. We used *“Classify Seed Point”* for the filter type in *“Quality”* section with *“Lower Threshold”* set at 40. We manually chose five representative holes during 24 min of observation as shown in Figure 3D.

### Autocorrelation function of the direction of collective migration

The direction of collective migration 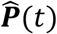 is defined from

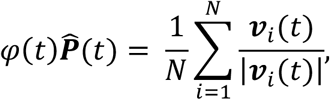

where *φ*(*t*) is the polar order parameter, *N* is the number of tracked cells, and ***ν**_i_*(*t*) is the instantaneous cell velocity of cell *i*. The autocorrelation function of the direction of collective migration 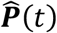 is given by

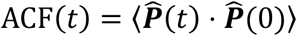

where 〈·〉 indicates the average over ensemble. The direction of collective migration 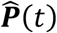 and its auto-correlation function ACF(t) were calculated for the cells in small regions of 125μm x 125μm along x- and y-coordinates, which is averaged for each sample.

### Measurement of aspect ratio

To obtain the aspect ratio of cell shape (Figure S5B), we rendered fluorescently labeled cell membrane using the “surface” function in IMARIS. The shortest length and the longest length were obtained from object-oriented Bounding Box OO statistical variables of IMARIS (Figure S5B, top right). The aspect ratio was then calculated by dividing the longest length by the shortest length (Figure S5B, bottom right).

### Theoretical Model for the formation of meshwork-like structure

In order to understand how the mesoderm cells organize into the meshwork structure, we introduce a mathematical model where each cell is represented by a self-propelled rod-shaped agent. To take into account the adhesion and volume exclusion between the cells, a short-range attractive interaction with a repulsive core is assumed between the agents. Since the typical size and migration speed of the cells are about 10 *μm* and 3 *μm/min*, we can assume that their dynamics is in the overdamped regime. Then, the equation of motion of the agent *i* is given by

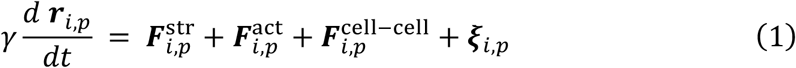

Here, the actual degrees of freedom for each rod-shaped agent are given by the head (*p* = 2) and tail (*p* = 1) particles of the diameter *d* that are separated by the length *ℓ_c_*, which gives the aspect ratio of the agent shape as *r* = (*ℓ_c_* + *d*)/*d*. ***r**_ip_* is the position of the tail and head particles of the agent *i*, and the friction coefficient *γ* = 3*πη*(3*d* + 2*ℓ_c_*)/5 takes into account the effect of the elongated shape with the effective viscosity *η*. The agent shape is kept the same by the stretching elasticity acting between the head and tail particles:

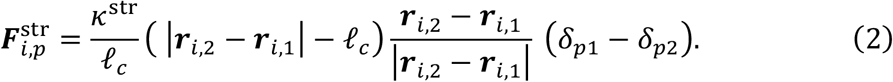

Here, *δ_pq_* is the Kronecker delta that takes 1 if *p* = *q* and 0 otherwise. A constant effective self-propulsion force of the magnitude *f^act^* is assumed acting only on the head particle as

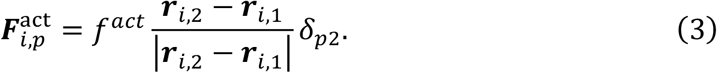

To implement the interaction between the agents, each rod-shaped agent is discretized into *M* helper particles, including the head and tail particles, of the equal distance less than 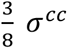, and the interaction force is imposed between the closest helper particles of a pair of agents (see below). The force on the pth helper particle is imposed on the head and tail particles with the geometric weight 1 – *α_p_* and *α_p_*, where *α_p_*|***r***_*i*,2_ – ***r***_*i*,1_| is the distance of the *p*th particle and the tail particle. As a result, the interaction force on particle *p* of agent *i* is given by

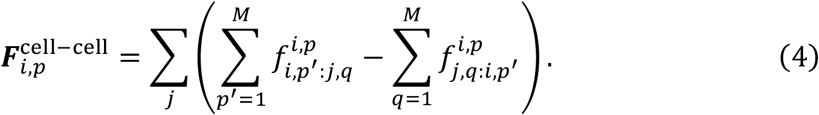

Here 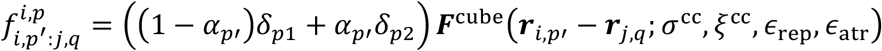, wtere *q* is the particle index of agent *j* that is the closest to particle *p*’ of agent *i*. Here, we use the following function of the short-range attraction with the repulsive core with the cutoff distance *r* < *σ*^cc^ + *ξ*^cc^ (55):

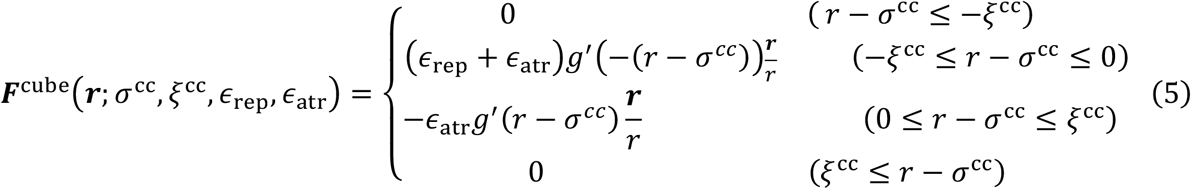

where 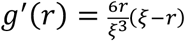 is the derivative of 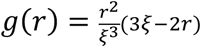. Finally, ***ξ***_i,k_ is a Gaussian white noise with zero mean and 〈*ξ_i,k,α_ ξ_j,l,β_*〉 = *σ δ_ij_δ_kl_δ_αβ_* with the noise strength *σ*.

To understand the essential aspect of the meshwork formation, we consider the model in a two-dimensional space in the range −*L_x_*/2 ≤ *x* ≤ *L_x_*/2, and −*L_y_*/2 ≤ *y* ≤ *L_y_*/2, where *L_x_* and *L_y_* are the system size. For the steady-state analysis, the periodic boundary conditions are assumed in both *x* and *y* directions. In the case where the cells are supplied from one *x* boundary in the manner as described below, the periodic boundary condition is assumed only in the *y* directions.

To mimic the experimental situation where the cells are supplied from the primitive streak, we prepared the source of agents at *x* = −*L_x_*/2 from which the agents are supplied at random y position at constant rate *r_source_*. In the source, the agents undergo random walk, without self-propulsion nor interaction with other agents, in a harmonic potential centered at *x_source_* = −(*L_x_* + *L_source_*)/2 that keeps the agents in the source. Here, *L_source_* = 2*ℓ_c_* is the width of the source. The agents that are supplied from the source experience the repulsive interaction from the source within the cutoff distance 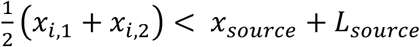,

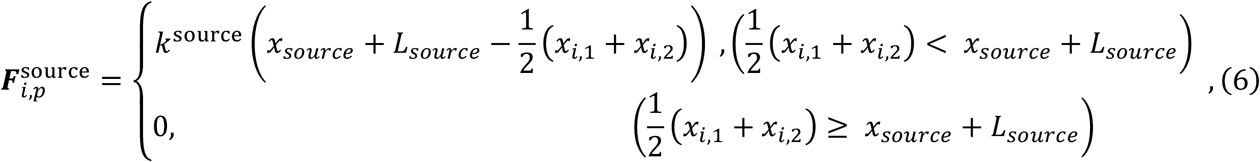

in addition to the self-propulsion and interaction force. Furthermore, in this case, we introduce the chemotactic force

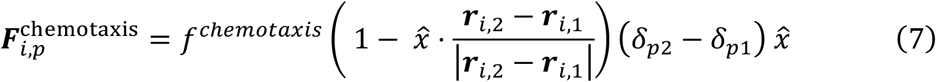

which rotates the agents so that they tend to move away from the source. The other x boundary is the sink of agents. That is, when the agents reach the boundary at *x* = *L_x_*/2, the agents are taken away from the system and placed back to the source. Therefore, the equation of motion of the agent *i* in this case is given by

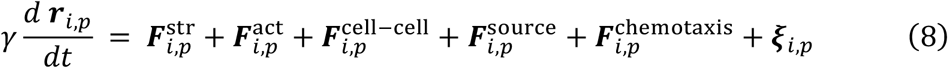

The parameters that were used in the numerical simulations are summarized in Table S6. In the case that the cells are supplied from the source, additional parameters are summarized in Table S7.

In the analysis shown in Figure 5F, to identify clusters of agents, we applied the Cluster analysis modifiers of Ovito Pro (56) to the simulation data including all the helper particles with the cutoff length *σ^cc^* + *ξ^cc^*/2. To quantify the elongation of the cluster, we measured the gyration tensor of each cluster defined by

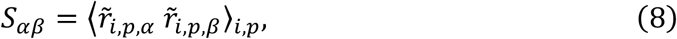

where 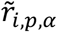 is the *α* component of the position of helper particle *p* of agent *i* measured from the center of the cluster, the average 〈·〉_*i,p*_ is calculated over helper particles *p* of all agents *i* that belong to the cluster. By using this gyration tensor, we calculated the cluster aspect ratio and the longitudinal angle as the square root of the ratio of the two eigenvalues, 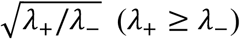, and as the direction of the major principal axis, respectively. To eliminate small clusters, we took into account only the clusters composed of more than four cells.

The nematic order and the nematic angle of the cells in a cluster shown in Figure 5F are calculated as the magnitude and angle of the nematic director defined by

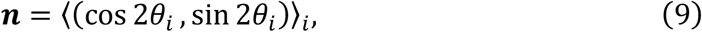

where *θ_i_* is the angle of the vector 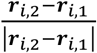 of cell *i*. The average 〈·〉_*i*_ is calculated over the cells that belong to the cluster.

### Correlation between the cluster elongation and nematic order

The correlation between the cluster elongation and the nematic order of the cells in the cluster (Figure 5F, right) is quantified by the order parameter defined by

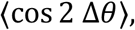

where Δ*θ* = *θ_l_* – *θ_n_* is the difference between the longitudinal angle *θ_l_* and the nematic angle *θ_n_* of each cluster. Here, note that both angles are of 2-fold rotational symmetry. To eliminate the effect of small or less-elongated clusters, the average 〈·〉_*i*_ is calculated over the clusters composed of more than four cells and the aspect ratio larger than or equal to 2.

---

**Figure S1 Motility analysis of mesoderm cells.** Related to Figure 1. (A) Analysis of the cell motility in the z-direction. Probability density function obtained from the six embryos indicates the frequency of the range of cell motion in z direction, which were obtained as the difference between the maximum and minimum z positions of individual trajectories that were in the image window for more than 60 min. The numbers of cells analyzed are N=1096, 350, 396, 302, 1455, 1044. The thick red line indicates the probability density function averaged over six samples. (B) Individual cell speed and progression speed in the anterior, middle, and posterior regions for the data shown in Figure 1D. The average (x) and the standard deviation (error bars) were shown. (C) Correlation between directionality and the MSD exponent. (D) Auto-correlation function of velocity for each sample.

**Figure S2. Meshwork structures in the mesoderm and quantitative analysis by persistent homology.** Related to Figure 2. (A) Transvers section of the embryo. (A1) Horizontal section of the whole chicken embryo at stage HH 4. (A2) Transverse section along the horizontal yellow line in (A1). (A3) Magnified view of the white box in (A2). The holes are marked by the yellow asterisks. Scale bars; (A1) 200μm; (A2) 50μm. (B) Correspondence of the points in the persistence diagram (PD) and the holes in the input image obtained by the advanced inverse analysis. The data corresponds to that in Figure 2E upper and 2F upper. (B1) The points with large lifetime in PD form a hole branch around death time ~ 0. (B2) Magnified view of the red box in B1 showing the hole branch of the points, which correspond to the holes in the input binary image (B3). The numbers assigned to each point in (B2) correspond to those in B3, which confirms the correspondence between the birth-death pairs and the holes. Scale bar, 30 μm in B2.

**Figure S3. 4D (*xyzt*) visualization of mesoderm cell migration using TG-GFP chick embryo.** Related to Figure 3. (A) Image of the time frame t=0 min of the live imaging. The transverse (horizontal) view was obtained at the level indicated by the yellow dotted line in the horizontal (transverse) view. Scale bars, 50 μm. See also Video S4. (B) Change of the meshwork structure over time. The holes in the images of 0 min (red), 15 min (yellow), 25 min (blue) and 35 min (green) are manually traced. (C) Displacement of the contour of the holes traced manually during 35 min. The holes move in the anterior-lateral direction during the observation as indicated by the arrows. Scale bar, 50 μm. (D) Input binarized images used for the persistent homology analysis and the extracted images by using the advanced inverse analysis. The five holes labelled by ①-⑤ are extracted to visualize their time evolution in Figure 3D by stacking them along the t axis.

**Figure S4 Statistical analysis of the collective migration of control cells and N-cadherin mutant expressing cells for the five embryos.** Related to Figure 4. Distribution of (A) individual cell speed, (B) directionality of individual cells, (C) progression speed of 50 μm x 50 μm areas, and (D) polar order parameter of 125μm x 125μm, and (E) autocorrelation function (ACF) of the direction of collective migration. (A)-(D)Each circle (o) plots the temporal average. The average (x) over (A-B) the cells and (C-D) the areas and their standard error (error bar) were shown. P-values indicated in the graphs were obtained from Wilcoxon rank sum test between the control and mutant cells. In embryo ID 1, 2 and 4, the polar order parameter on the control side was higher than that on the mutant side. In embryo ID 3 and 5, the difference was not statistically significant. (E) The average and the standard error of means of ACF calculated for each subarea (125μm x 125μm) are shown.

**Figure S5. Over-expression of N-Cadherin mutant changes in cell morphology and intercellular properties.** Related to Figures 4 and 5. (A) Snapshots of the live imaging of the control cells (upper panels) and the N-Cad-M expressing cells (lower panels). Cell membrane is marked by GFP-CAAX. The numbers (1-5 in the images of control cells, 1-7 in the images of N-cadherin mutant cells) are assigned to track the cells. Scale bars, 10 μm. (B) Multiphoton images of the control cells and the N-Cad-M expressing cells (“Raw image” in the left panel). Cell membrane is marked by GFP-CAAX and extracted by surface function of IMARIS software (“Extracted cells by Surface Function” in the left panel). (B right top panel) A schematic diagram shows the shortest length and the longest length of a cell. a: Length of the shortest principal axis. b: Length of the longest principal axis. The software identifies a cell by considering the object-oriented minimal rectangular box, as shown in red. (B right bottom panel) Aspect ratio of the control cells and the N-Cad-M expressing cells. Error bars are the standard error of mean; N=51 (control), N=37 (N-cad mutant). Asterisk, p < 0.001 (t-test).

**Video S1. Mesoderm cell movements on gastrulating chick embryo.** Related to Figure 1. Left: Nuclei of mesoderm cells are labelled by H2B-eGFP expression (green). Right: Cell trajectories by IMARIS tracking in the anterior, middle and posterior regions. Scale bars, 50 μm.

**Video S2. Meshwork structure in the mesoderm.** Related to Figure 2. Confocal Z-stack images of the mesoderm and the primitive streak of stage HH4 chick embryo. The embryo is stained for nuclei with DAPI (cyan) and for N-cadhein (green). Z-stack images with a thickness of 1.5 μm show that the characteristic meshwork structures are composed of multiple cells in the mesoderm located on both sides of the primitive streak. ps, primitive streak.

**Video S3. Mesoderm cells forming a dynamic meshwork during migration.** Related to Figure 3. Live imaging of a thin section (5μm) of the mesoderm of stage HH4 GFP expressing transgenic chicken embryo. The mesoderm cells migrate from the primitive streak by forming a dynamic meshwork structure undergoing continual and rapid re-organization. ps, primitive streak. Scale bar, 50μm.

**Video S4. Dynamics of the meshwork structure.** Related to Figures S3A and B. 4D live imaging of the mesoderm of stage HH4 GFP expressing transgenic chicken embryo. The optical transvers section and horizontal section, monitored for 40 min, shows that the three-dimensional dynamic meshwork structure is formed by the migrating mesoderm cells and the holes move toward the anterior-lateral direction. ps, primitive streak. Scale bar, 50 μm.

**Video S5. Cell trajectories of the N-Cadherin mutant expressing cells and the control cells.** Related to Figure 4. Left, Control side: Nuclei of mesoderm cells are labelled by H2B-eGFP expression (control, green). Most mesoderm cells away from the primitive streak migrate toward the anterior-lateral direction. Right, N-cadherin mutant side: The N-cadherin deletion mutant expressing cells are detectable by H2B-mCherry expression. These cells also migrate in the anterior-lateral direction but exhibit zigzag trajectories, which is apparently different from the control cells. Scale bars, 30μm.

**Video S6. Cell-cell contact behaviors in the N-Cadherin mutant expressing cells**

**and the control cells.** Related to Figure S5A. Cell membrane of the control cells and of the N-Cad-M expressing cells are detected by GFP-CAAX expression. Left: The control cells undergo continual contact with the surrounding cells. Right: The N-Cad-M expressing cells change the contact partners one after another during the observation. Scale bars, 10 μm.

**Video S7. Meshwork structure formation in the simulation with agent supply.**

Related to Figure 5H. The agents were supplied from the PS boundary (left boundary). The head particle and the tail particles of an agent are indicated by blue and magenta colors, respectively.

**Table S1.** Experimental Models: Organisms/Strains

|  |  |
| --- | --- |
| Wild Type Chicken Eggs | Shimojima farm,<br>Kanagawa, JP |
| Tg (pLSi/ΔAeGFP) Chicken Eggs | ABRC, University<br>of Nagoya; Motono<br>et al. 2010 |

**Table S2.**
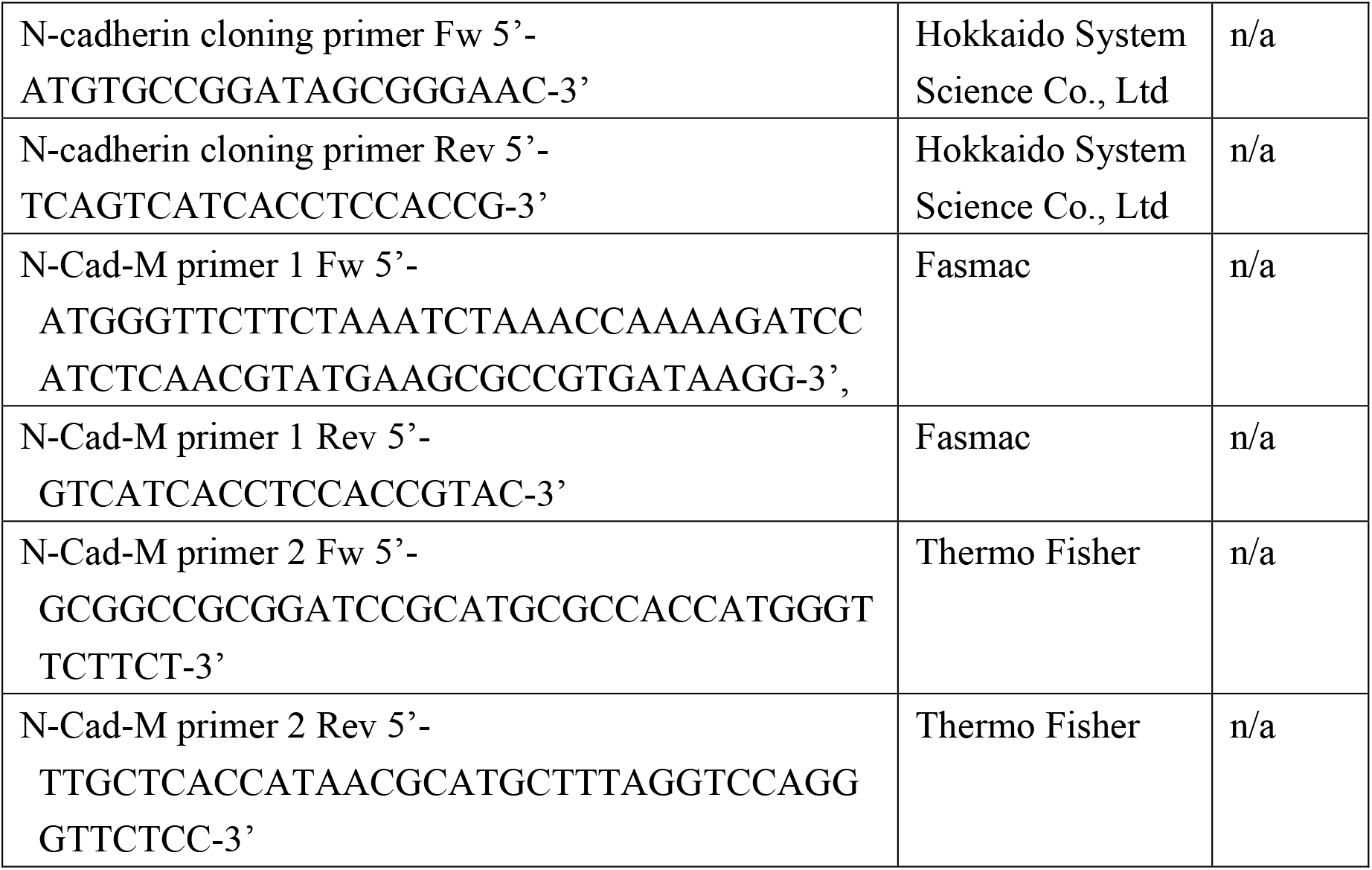
Oligonucleotides used in this study

**Table S3.** Recombinant DNA constructed in this study

|  |  |  |
| --- | --- | --- |
| pGEM-T Easy | Promega | Cat# A1360 |
| pCAG-H2B-eGFP | Dr. Hadjantonakis<br>(MSKCC, NY). |  |
| pCAG-N-Cad-M-2A-H2B-mCherry | This study |  |
| pCAG-N-Cad-M-2A-eGFP-CAAX-2A-H2B-mCherry | This study |  |

**Table S4.** Antibodies and chemicals used in this study

| Antibodies | Source | IDENTIFIER |
| --- | --- | --- |
| Purified Mouse Anti-E-Cadherin<br>1:1000 dilution | BD transduction<br>Lab | Cat# 610181; RRID:<br>AB_397580 |
| Anti-N-cadherin, polyclonal<br>1:300 dilution | Takara bio | Cat# M142; RRID: AB_444317 |
| Monoclonal Anti-N-Cadherin/A-41CAM (Clone GC-4)<br>1:50 dilution | Sigma-Aldrich | Cat# C2542; RRID: AB_258801 |
| Goat anti-Mouse IgG, (H+L)<br>Highly Cross-adsorbed<br>Antibody, Alexa Fluor 488<br>1:300 dilution | Thermo Fisher | Cat# A-11029; RRID:<br>AB_138404 |
| Goat anti-Rabbit IgG, (H+L)<br>Highly Cross-Adsorbed<br>Secondary Antibody, Alexa<br>Fluor 488<br>1:300 dilution | Thermo Fisher | Cat# A-11034 ;<br>RRID:AB_2576217 |
| Cellstain DAPI Solution<br>1:100 dilution | Dojindo<br>Laboratories | Cat# D523 |

**Table S5.** Software and Algorithms used in this study.

|  |  |  |
| --- | --- | --- |
| Fiji | NIH | https://fiji.sc |
| Imaris x64 9.5.1 | Oxford Instruments | https://imaris.oxinst.com/<br>RRID:SCR_007370 |
| HomCloud | HomCloud<br>Development team | https://homcloud.dev/index.en.html |
| Custom MATLAB<br>code for analyzing<br>cell trajectories | This study | N/A |
| Custom C++ code for numerical simulation | This study | N/A |
| Ovito Pro |  | https://www.ovito.org |

**Table S6.** The parameters used in the numerical simulation.

| *variables* | *symbols* | *values* |
| --- | --- | --- |
| *Cell diameter* | $d$ | 1 |
| *Cell head and tail length* | $\ell_c$ | $d(\alpha - 1)$ |
| *Cell stretching elasticity* | $\kappa^{str}$ | 10 |
| *Cell-cell attraction strength* | $\epsilon^{atr}$ | 0.01 |
| *Cell-cell repulsion strength* | $\epsilon^{rep}$ | 0.1 |
| *Cell-cell interaction length* | $\sigma^{cc}$ | $d$ |
| *Width of cell-cell attraction well* | $\xi^{cc}$ | $d/2$ |
| *Effective viscosity* | $\eta$ | 0.1 |
| *Temperature* | $k_B T$ | 0.004142 |
| *Noise strength* | $\sigma$ | $2 k_B T \gamma$ |
| *Number of cells* | $N_{cell}$ | 1600 |

**Table S7.** The parameters used in the numerical simulation with the agent supply.

| *variables* | *symbols* | *values* |
| --- | --- | --- |
| *Self-propulsion force* | $f^{act}$ | 0.01 |
| *Repulsive strength from source* | $k^{source}$ | 0.1 |
| *Chemotactic force* | $f^{chamotaxis}$ | 0.004 |
| *Simulation box size in $x$* | $L_x$ | 60 |
| *Simulation box size in $y$* | $L_y$ | 40 |

## Notes

### Competing Interest Statement

The authors have declared no competing interest.

