## Supplementary figures and images for "Migrating mesoderm cells self-organize into a dynamic meshwork structure during chick gastrulation"

### Supplemental Figures

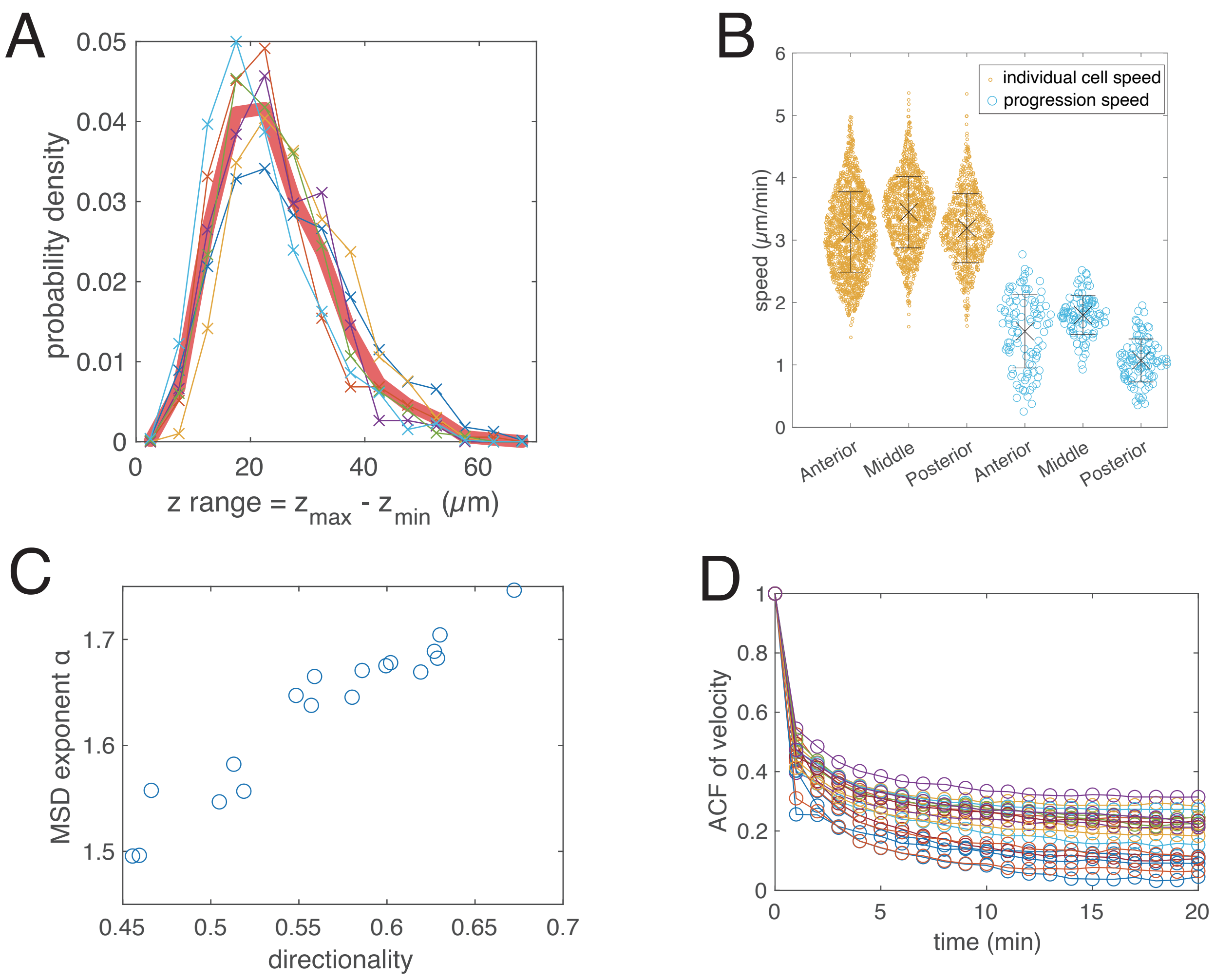

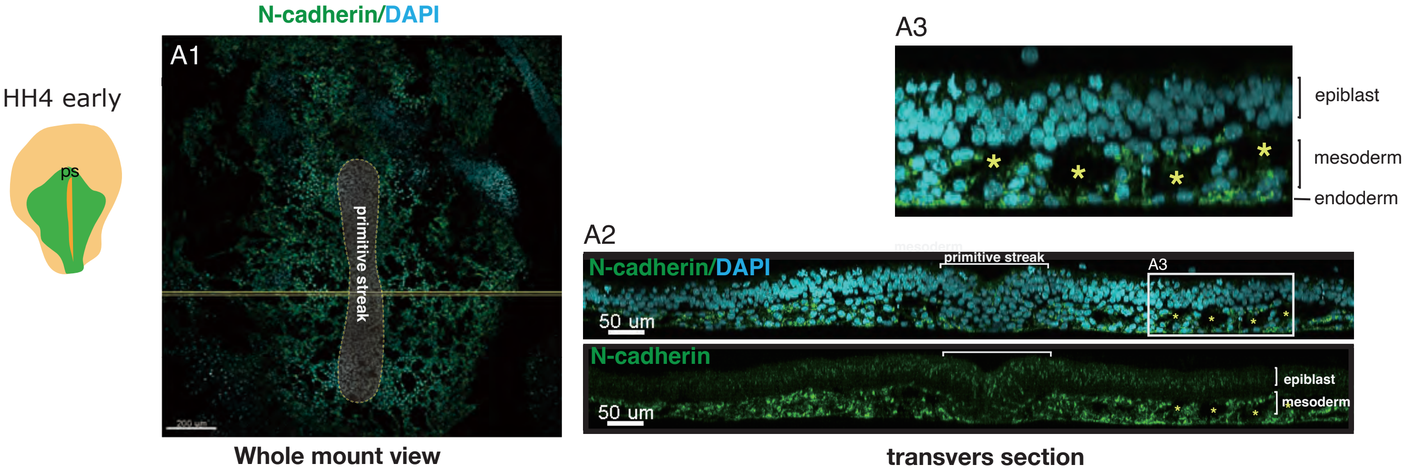

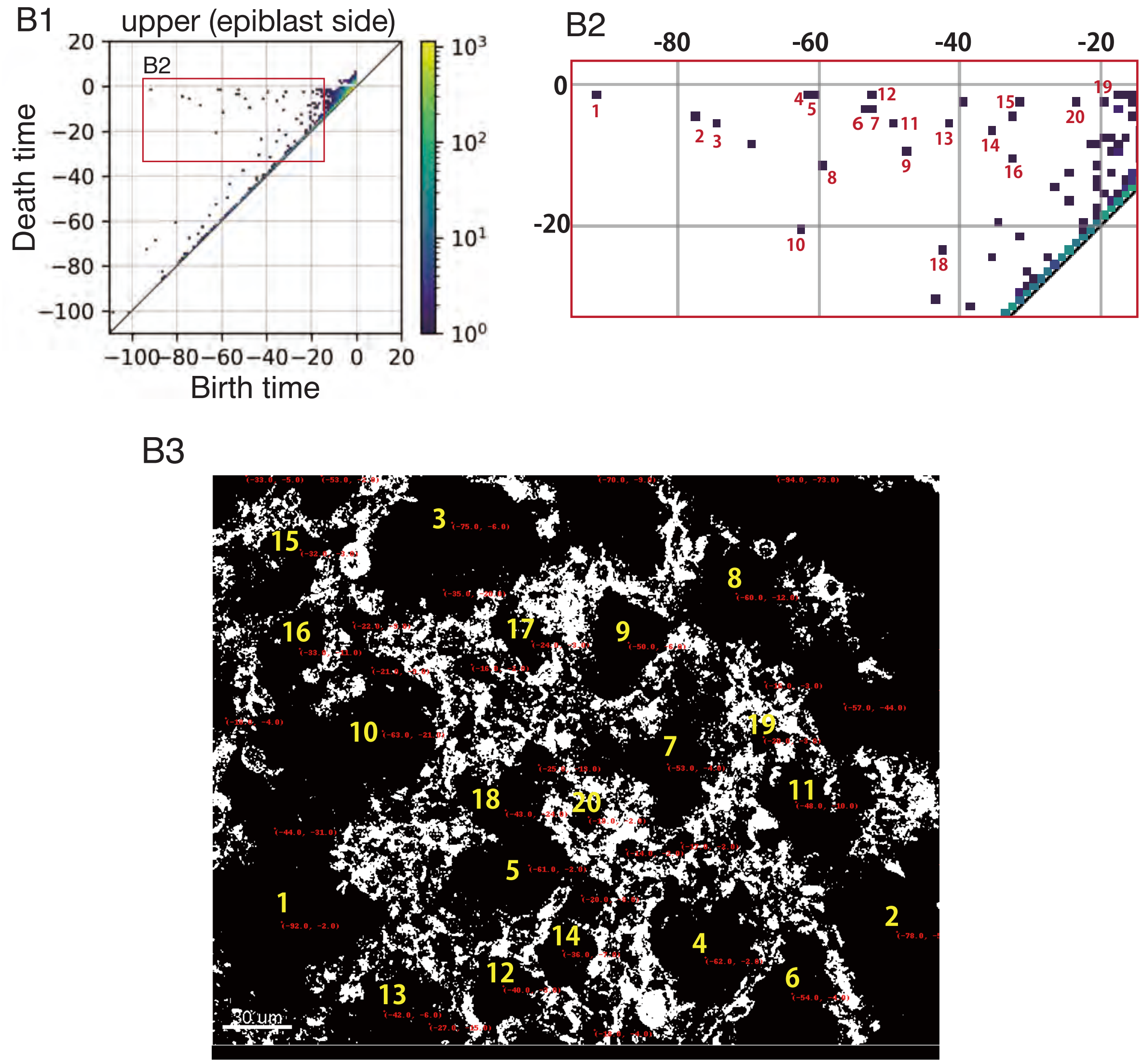

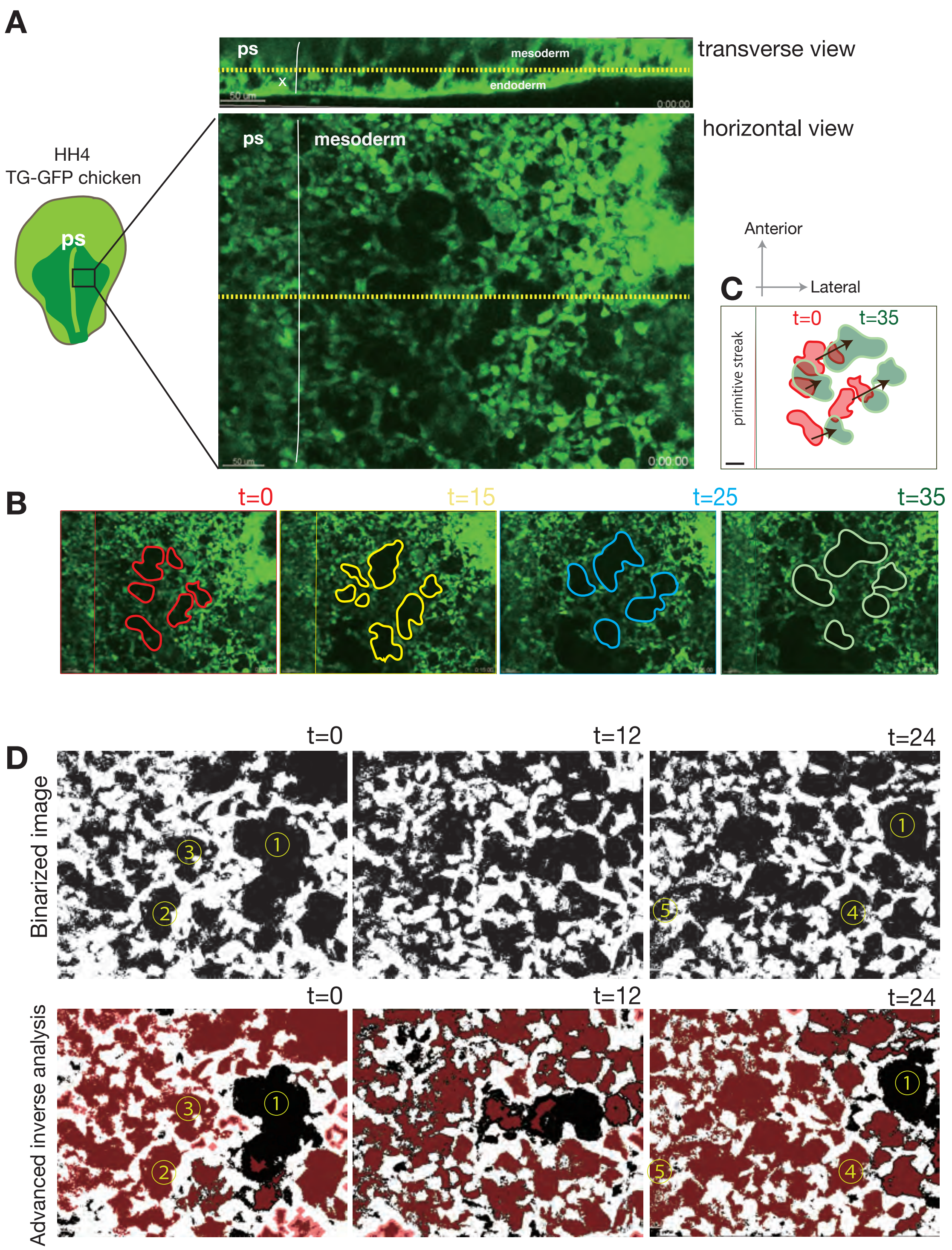

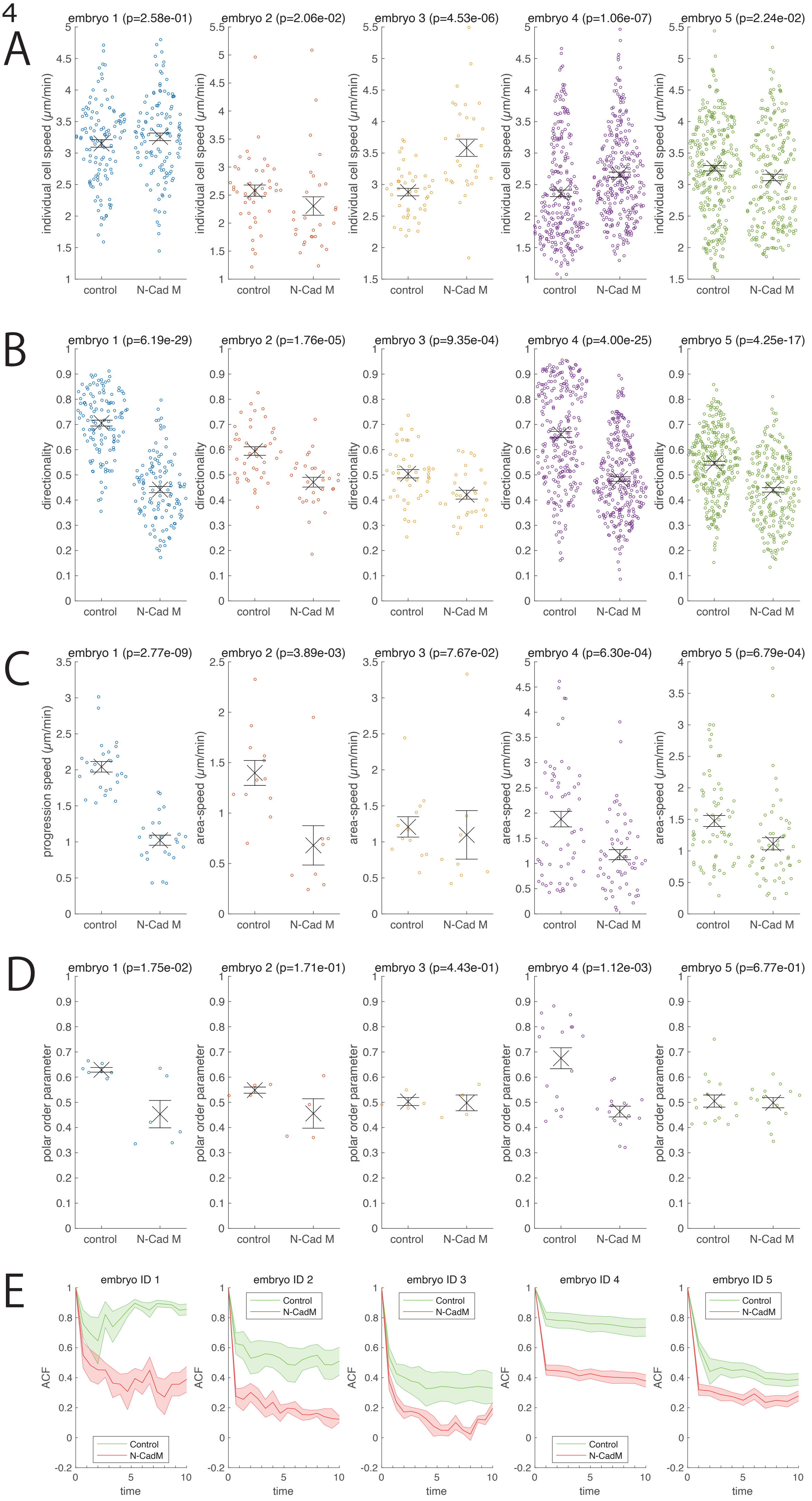

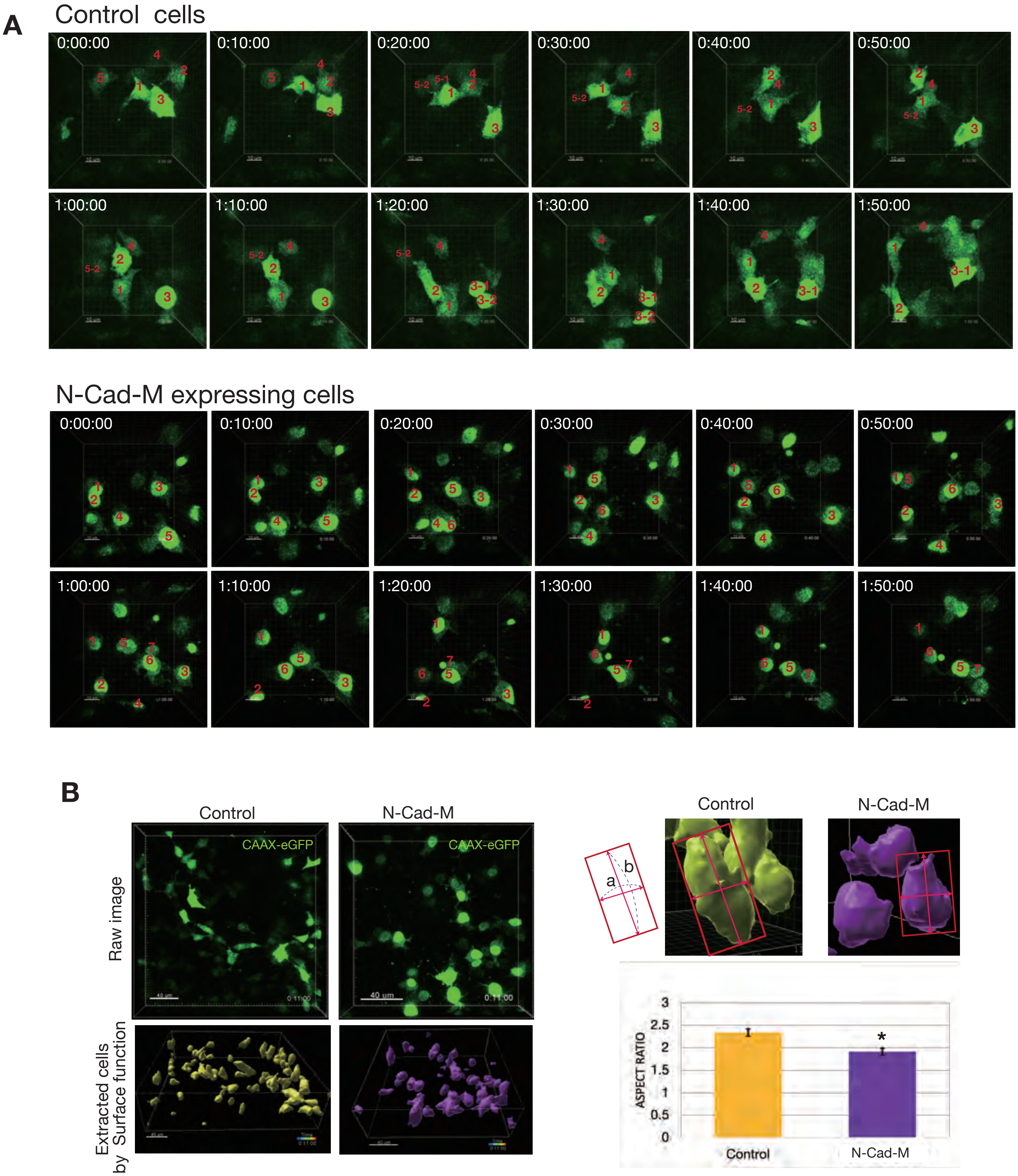
